# Multidomain Coupling Governs FoxP1 Assembly and Nuclear Compartmentalization

**DOI:** 10.64898/2026.06.02.728184

**Authors:** Anay F. Lazaro-Alfaro, Javiera Avilés, Thomas-Otavio Peulen, Exequiel A. Medina, Katrin G. Heinze, Hugo Sanabria, Katherina Hemmen

## Abstract

FoxP1 is a multidomain transcription factor implicated in development, immunity, and cancer, widely proposed to function as a dimer. However, the molecular mechanisms governing its assembly and nuclear organization in living cells remain unclear.

Here, we combine biochemical assays and live-cell fluorescence lifetime imaging to resolve FoxP1 homotypic interactions. We show that FoxP1 forms heterogeneous complexes whose stability is governed by antagonistic coupling between its leucine-zipper (ZIP) and Forkhead (FKH) domains. The FoxP1 ZIP domain promotes dimerization while suppressing FKH-mediated interactions, revealing a competing interdomain mechanism that tunes complex formation. Pathogenic and deletion variants disrupt this intricate balance, altering interaction stability and promoting the formation of dense nuclear condensates upon loss of DNA binding. Together, our results demonstrate that FoxP1 assembly is encoded by its multidomain architecture. Our findings show how competing interaction domains regulate transcription factor complex formation, nuclear organization, and DNA binding, with implications for disease-associated dysregulation.

**GRAPHICAL ABSTRACT:** 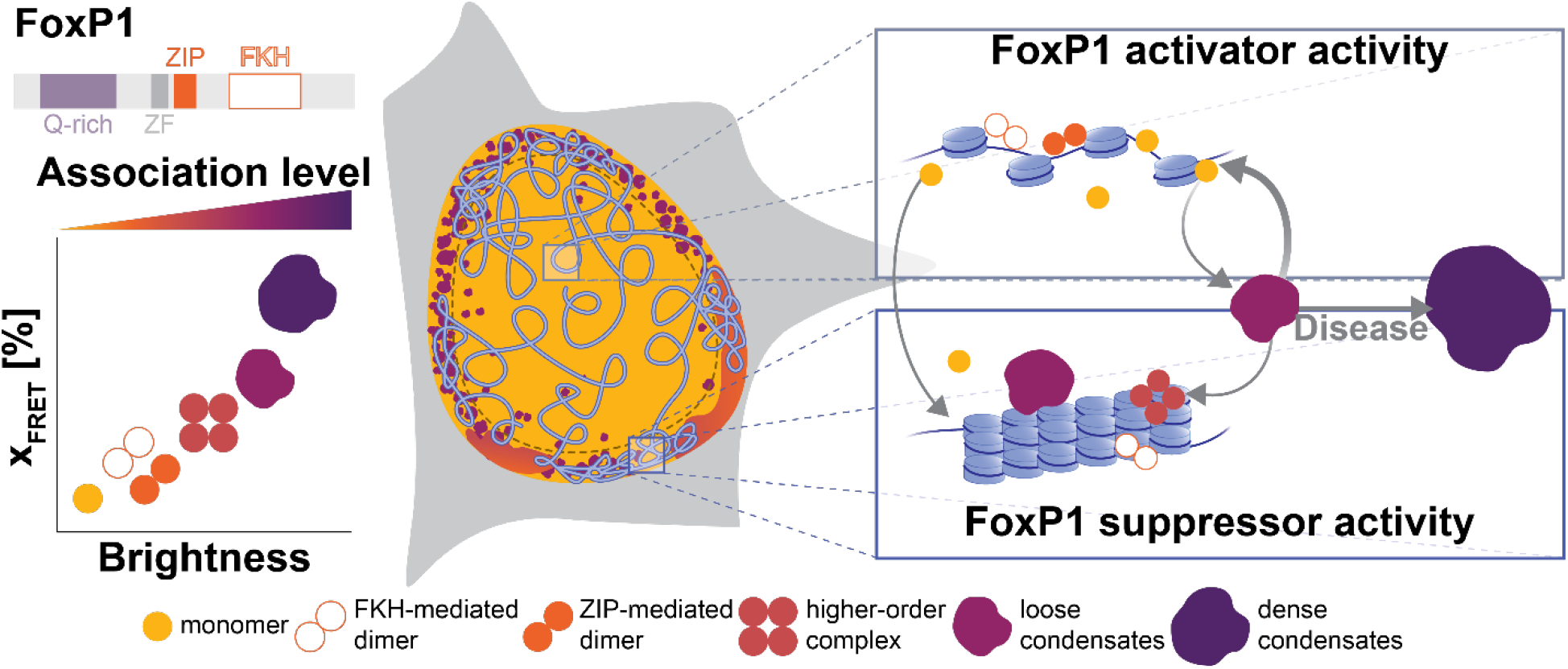

**HIGHLIGHTS:**

- FoxP1 complexes assemble through two distinct mechanisms mediated by the FKH and ZIP domains.
- Interdomain coupling switches the configuration of the FoxP1 complex.
- FKH-mediated FoxP1 oligomers and other higher-order complexes accumulate at the nuclear periphery, whereas ZIP-mediated dimers remain more uniformly distributed.
- Disruption of FKH-DNA binding drives formation of protein-dense FoxP1 condensates.

## INTRODUCTION

A precise regulation of gene expression is essential for maintaining cellular identity and proper physiological function. Transcriptional output arises from coordinated interactions among chromatin architecture, transcription factors (TFs), and the basal transcriptional machinery. TFs bind specific DNA sequences to regulate gene accessibility and drive dynamic changes in genome organization^1-3^. These proteins contain conserved DNA-binding domains that confer high target specificity^4,5^ and define major TF families, including Forkhead-box (Fox) proteins^6^.

Beyond their DNA-binding domains, TFs frequently contain structural domains that regulate protein-protein interactions, thereby influencing transcriptional output. These include oligomerization domains such as leucine zippers and zinc fingers^7-12^. In many multidomain TFs, dimerization represents a prevalent regulatory strategy that enhances DNA-binding affinity, sequence specificity, and overall regulatory versatility^6,13,14^.

Notably, more than half of eukaryotic TFs contain extensive intrinsically disordered regions (IDRs) or low-complexity domains^15-18^, which promote molecular flexibility and condensate formation^19-22^. However, these regions complicate structural characterization. Consequently, the cooperativity of distinct domains in the formation of functional TF complexes remains poorly understood. The multivalence of these multidomain transcription factors suggests a regulation through coordinated interactions between structured domains and disordered regions.

Here, we focus on FoxP1, a member of the P-subfamily of Fox transcription factors, to investigate the regulatory role of multidomain architecture in homotypic protein complex formation. Fox proteins are evolutionarily conserved regulators^23^ involved in development, immunity, and cancer^24-26^. They share a conserved Forkhead (FKH) domain that mediates DNA-binding^27^. Members of the P-subfamily additionally contain a leucine zipper (ZIP) domain that promotes dimerization^28-30^, and a C2H2-like zinc finger (ZF) domain implicated in regulatory interactions^31^. FoxP1 also harbors a glutamine-rich, low-complexity region (Q-rich), whose functional role remains largely unexplored (**Figure 1A**).

**Figure 1.**
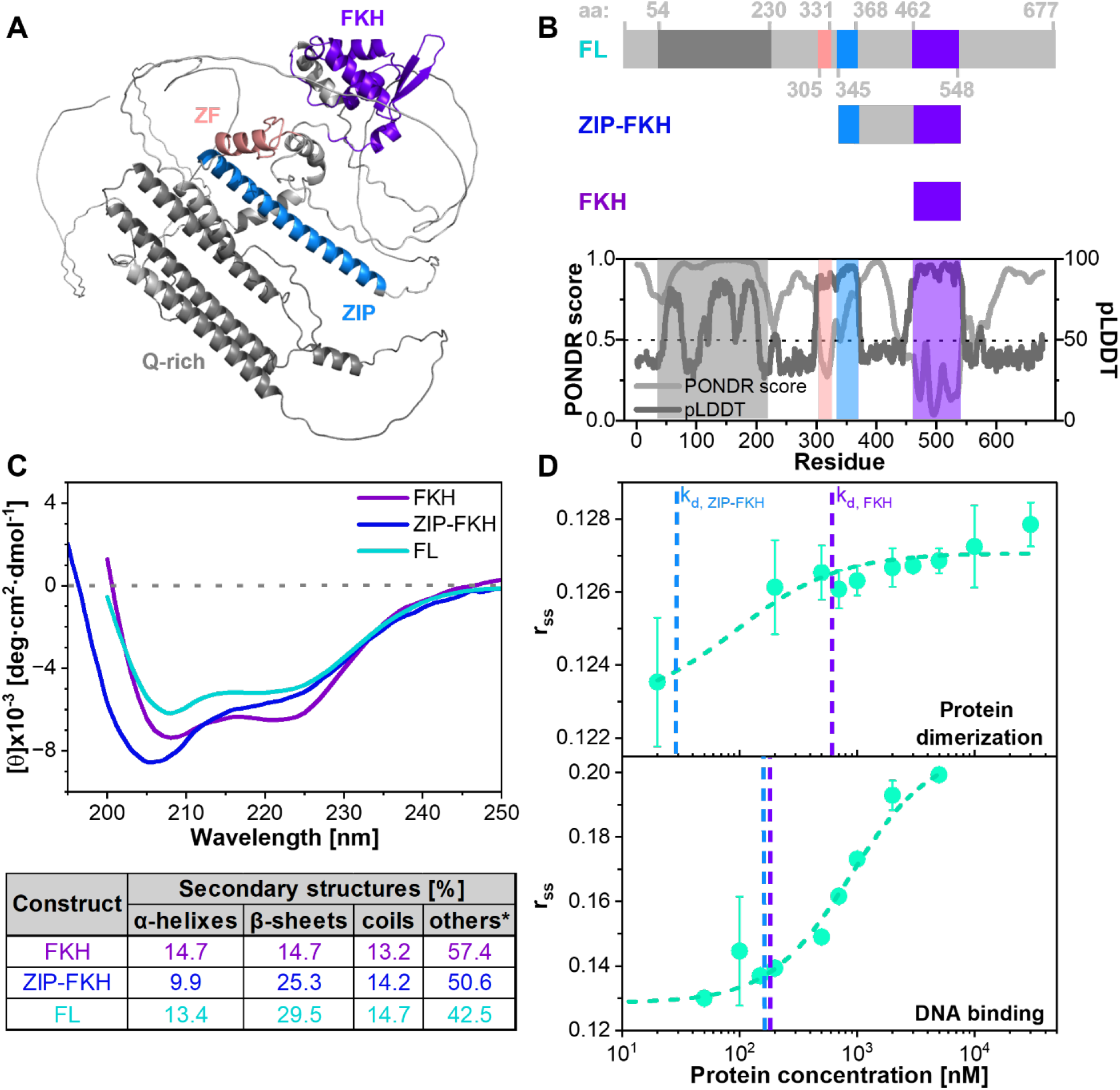
Full-length FoxP1 displays unexpected structural organization that modulates dimerization and DNA binding. (**A**) AlphaFold2^47,48^ prediction of full-length (FL) FoxP1 structure (average pLDDT 57.9%), showing the glutamine-rich region (Q-rich, dark grey), zinc finger domain (ZF, salmon), leucine zipper domain (ZIP, blue) and forkhead domain (FKH, purple). (**B**) Linear representation of FL FoxP1 and truncated fragments (ZIP-FKH and FKH) used for *in vitro* studies, color-coded as described in (A). The lower panel shows the Predictor of Naturally Disordered Regions (PONDR, light grey) score^49^ and predicted Local Distance Difference Test score (pLDDT, dark grey) from the AlphaFold model. (**C**) Circular Dichroism spectra of FL, ZIP-FKH, and FKH constructs. The table shows the secondary structure content of each construct estimated using BeStSel^51,52^. *Others include: 3_10_-helix, π-helix, β-bridge, bend, loop/irregular and invisible regions. (**D**) Steady-state fluorescence anisotropy (*r*_*ss*_) measurements were used to analyze FoxP1 dimerization (upper panel) using 20 nM labeled FL FoxP1, and DNA binding (lower panel) with a fixed DNA concentration (180 nM unlabeled + 20 nM labeled, Materials and Methods). In both assays, increasing concentrations of unlabeled FL FoxP1 were titrated, and the dissociation constants (K_d_) were estimated as 93 ±77 nM for dimerization and 879 ±104 nM for DNA-binding. Blue and purple dotted lines indicate previously reported K_d_ for the ZIP–FKH and FKH fragments, respectively ^28,37^. These results reveal that the structural context of full-length FoxP1 alters both its oligomerization behavior and DNA-binding affinity relative to truncated fragments.

Early *in vitro* experiments showed that FoxP1 binds DNA as a ZIP-mediated dimer^32^. However, its modular architecture suggests the existence of additional interaction interfaces. Structural studies later demonstrated that FoxP1 FKH domain can form weakly stable domain-swapped dimers^27,33-37^. Importantly, DNA binding destabilizes this domain-swapped state^38^, raising questions about its biological relevance. Moreover, while the ZIP domain promotes dimerization^8^, it suppresses FKH domain swapping in FoxP^28^, suggesting the presence of distinct and potentially competing dimerization mechanisms in isolated protein. These findings suggest that antagonistic interdomain coupling is a potential regulatory mechanism of FoxP1 complex formation. Consistent with that idea, FoxP1 has been reported to function both as a transcriptional repressor^39,40^ and as a regulator that enhances protein expression^41^, suggesting that it forms multiple functional complexes.

Despite these advances, the structural organization and functional *in situ* assembly of full-length (FL) FoxP1 remain unknown. This gap is especially relevant given FoxP1’s proposed pioneer-like activity^42-46^, which requires coordinated interactions with chromatin and regulatory machinery. Here, we defined the DNA-binding mechanism of FoxP1 as the integrated arrangement of domain orientation, conformational state, and interaction interfaces within functional dimers or higher-order oligomers. We hypothesize that FoxP1 regulates gene expression by modulating its assembly into distinct states in a manner sensitive to nuclear context.

To test this hypothesis, we combine computational analysis, biochemical assays in isolated proteins, and live-cell fluorescence lifetime imaging to address two central questions: (i) how FoxP1 domains cooperate to mediate FoxP1 homotypic protein interactions, and (ii) how distinct homo-complex states correlate with nuclear organization. By elucidating the organizational principles governing FoxP1 homo-complex assembly, this work establishes a framework for understanding how multidomain transcription factors regulate gene expression in development and disease.

## RESULTS

### Full-length FoxP1 displays unexpected structural order and reduced DNA binding affinity

As a transcription factor, FoxP1 coordinates homotypic interactions and DNA binding within the structurally diverse chromatin environments of the nucleus. Understanding how FoxP1 domain organization supports gene regulation requires defining its conformational landscape and determining how this landscape remodels upon binding to distinct chromatin structures. Notably, sequence analysis predicts that FoxP1 contains IDRs, suggesting a high degree of structural plasticity, which complicates the elucidation of its conformational organization.

To gain initial insights into structural features that may inform FoxP1 quaternary organization, we examined the AlphaFold2-predicted tertiary structure of isolated FoxP1 retrieved from the AlphaFold Protein Structure Database^47,48^ (**Figure 1A**). The model yielded an average predicted Local Distance Difference Test (pLDDT) score of 57.9%, consistent with FoxP1’s high intrinsic disorder content, indicating that structured regions are largely confined to the folded domains and suggesting substantial conformational plasticity across the remainder of the sequence (**Figure 1B**, lower). Consistently, calculations of the VSL2 Predictor of Naturally Disordered Regions (PONDR)^49^ revealed that ∼79% of the sequence exceeds the disorder threshold (0.5), with structured regions largely confined to the ZF and FKH domains (**Figure 1B**). Interestingly, the two computational approaches differed in their assessment of the Q-rich region, suggesting the presence of collapsed IDRs, as discussed below.

To experimentally estimate FoxP1’s global secondary structure, we isolated FoxP1 fragments comprising the ZIP and FKH domains (ZIP-FKH, residues 345-548) or the FKH domain alone (FKH fragment, residues 462-548) (**Figure 1B**, upper) and compared their circular dichroism (CD) spectra with the FL FoxP1 spectrum (**Figure 1C**, upper). The FKH fragment displayed the characteristic minima at ∼208 and ∼222 nm, consistent with a predominantly α-helical secondary structure; notably, the deeper minimum at 208 nm relative to 222 nm indicates a substantial contribution from disordered or flexible regions^50^. Consistent with this interpretation, secondary structure estimation using BeStSel^51,52^ (**Figure 1C**, bottom) classified the FKH fragment as belonging to the α/β protein class, in agreement with its experimentally determined three-dimensional structure (PDB ID: 2KIU^34^).

Interestingly, the ZIP-FKH fragment exhibited an elevated β-sheet content (25.3%), which further increased to 29.5% in the FL construct. This estimation contrasts with predictions for FL FoxP1 derived from AlphaFold (**Figure 1A**), which showed mainly α-helical structures alongside unfolded regions. Notably, the extracted information from FL’s CD spectrum also exhibited the lowest proportion of disordered regions (42.5%) among all constructs analyzed, in contrast with the calculated PONDR score and closer to the AlphaFold prediction, suggesting that additional regions of the protein contribute to global secondary structural organization. The lower disordered fraction estimated by CD relative to PONDR, together with the pLDDT score mentioned above, likely reflects the presence of collapsed IDRs within the Q-rich region. In some cases, collapsed IDRs adopt helical or extended secondary structure detectable by CD, but lack the stable tertiary contacts required for high PONDR confidence scores^49^. This distinction is important: collapsed IDRs retain conformational flexibility while contributing to global secondary structure content, and may represent functionally relevant interaction-competent regions within the disordered landscape of FoxP1, as observed in other TFs^53,54^.

These structural differences prompted us to examine how the full-length architecture influences FoxP1 dimerization and DNA-binding affinity using fluorescence anisotropy measurements (**Figure 1D**). In fluorescence anisotropy experiments, increases in anisotropy (**Equation 1**) reflect increases in the effective hydrodynamic volume of the labeled molecule and, in this context, are related to protein association. To evaluate FoxP1 homodimerization, we titrated unlabeled FL FoxP1 into a solution containing fluorescently labeled FL FoxP1 (Materials and Methods). Unlike the ZIP-FKH and FKH fragments^28,37^, the addition of ∼100 nM unlabeled FL protein was sufficient to induce a measurable anisotropy transition, suggesting formation of a protein complex. Fitting the association curve to a single-binding model yielded a dissociation constant, K_d_, of 93 ±77 nM. This value falls between those previously reported for ZIP-FKH dimers^28^ and the isolated FKH domain^37^ (**Figure 1D**, top), indicating an intermediate association strength relative to the truncated constructs. These results suggest that either the C-terminal, the Q-rich region, or both contribute to FoxP1 complex formation.

We next examined DNA binding by FL FoxP1 using an analogous fluorescence anisotropy assay, where increasing concentrations of unlabeled FL FoxP1 were titrated into a mixture containing unlabeled DNA and fluorescently labeled DNA (**Figure 1D**, bottom). Fitting the binding isotherm yielded a K_d_ of 879 ±104 nM for the FL FoxP1-DNA interaction. Strikingly, this value was fivefold higher than that previously observed for the monomeric FKH and ZIP-FKH fragments^28^, indicating that the additional regions present in the full-length protein – including the Q-rich and the C-terminal region – modulate FoxP1 DNA binding. Together, these results suggest that the structural context of FL FoxP1 reshapes both protein-protein and protein-DNA binding behavior, highlighting the contribution of regions outside the canonical DNA-binding domain.

### FoxP1 variants form homotypic complexes in HEK293T nuclei

Because protein behavior often depends on cellular context, we examined whether the properties of FL FoxP1 observed in isolated proteins are maintained in the nuclear environment of living cells. To this end, we transiently transfected HEK293T cells with fluorescently tagged FoxP1 constructs. These included wild-type (WT) FL FoxP1 as well as truncated ZIP–FKH (residues 345–677) and FKH (residues 462–677) fragments, each retaining the C-terminal region (residues 549–677), which was absent in the constructs used for the *in vitro* assays. To ensure nuclear translocation of the truncated proteins, the FoxP1 nuclear localization sequence (NLS) (RRRYSD) was included at the N-terminus of all constructs. Constructs were fused C-terminally to either eGFP or mCherry, as N-terminal tags sequestered nuclear translocation (**Supplementary table 1**), even with the inclusion of the NLS.

Confocal imaging confirmed nuclear localization of FL FoxP1 and its fragments by counterstaining the nuclei with Hoechst dye to visualize DNA (**Figure 2A, Supplementary Figure 1**). FoxP1 fluorescence was enriched at the nuclear periphery, a region typically associated with transcriptionally inactive heterochromatin^55^. Notably, FL FoxP1 formed punctate structures within this region, whereas the fluorescence in the nuclear center appeared more diffuse. Line profiles across selected regions of interest (ROIs) revealed coincident intensity peaks in the eGFP/mCherry and Hoechst channels, indicating local enrichment of FoxP1 in DNA-rich regions

**Figure 2.**
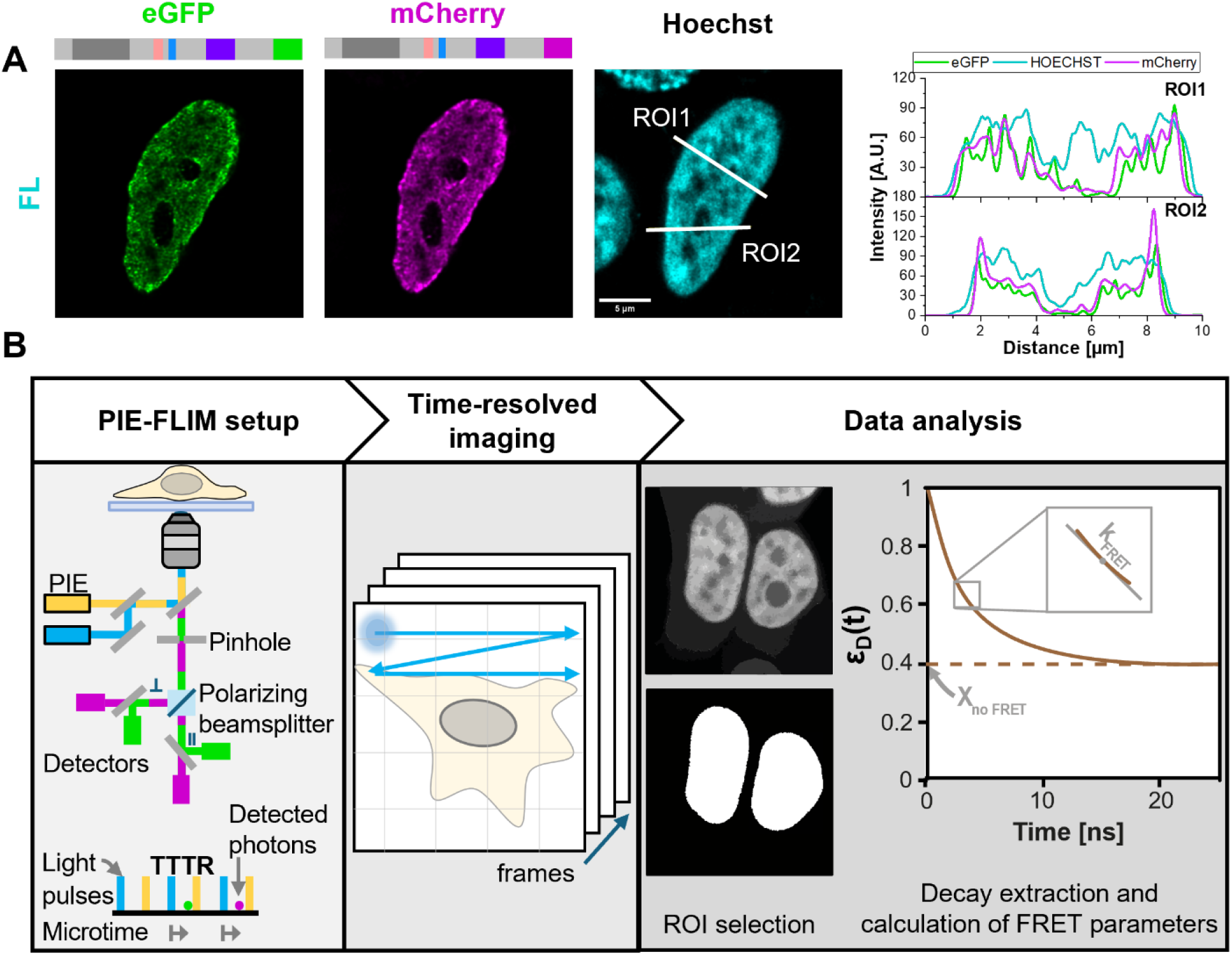
FoxP1 forms nuclear complexes enabling PIE-FLIM-FRET analysis of homotypic interactions. (**A**) Confocal image of a Hoechst-stained HEK293T cell nucleus co-transfected with eGFP-and mCherry-tagged full-length FoxP1 constructs, showing nuclear expression of both constructs. Line profiles of the two regions of interest (ROIs) indicate fluorescence intensity across the three detection channels. (**B**) Principle of confocal polarization-resolved PIE-FLIM-FRET measurements: Donor (green, 485 nm laser) and acceptor (magenta, 561 nm laser) excitation pulses are alternated during image scanning. In time-tagged time-resolved (TTTR) detection mode, the arrival times of each photon relative to the preceding laser pulse (microtime) are recorded together with the detection channel (green/magenta, parallel/perpendicular) and image frame markers. Selected nuclei or subnuclear ROIs are used to extract fluorescence intensity decays from which FRET parameters are calculated.

To quantify FoxP1 homotypic interactions within the nuclear environment, we used pulsed interleaved excitation fluorescence lifetime imaging microscopy (PIE-FLIM)^56,57^combined with Förster resonance energy transfer (FRET)^58^ (**Figure 2B**). In this approach, eGFP and mCherry serve as donor and acceptor fluorescent proteins, respectively, and are directly excited alternately on the nanosecond timescale. PIE defines two detection time windows: the prompt region, corresponding to fluorescence emission following donor direct excitation, and the delay region, corresponding to fluorescence emission following acceptor direct excitation. Energy transfer between fluorescent proteins reduces the donor fluorescence in the prompt region and its lifetime, providing quantitative readouts of protein proximity and complex formation in living cells.

Fluorescence photons were recorded in a time-tagged time-resolved (TTTR) format, with their arrival times relative to the nearest donor-excitation pulse (microtime). This measurement mode allows reconstruction of fluorescence lifetime images and intensity maps at pixel resolution. This analysis enabled us to define nuclei as regions of interest and to extract fluorescence intensity decay curves from them. Inspection of these decays provided qualitative indications of energy transfer, while quantitative analysis of fluorescence lifetimes and decays allowed us to map subnuclear heterogeneities and assess the contribution of individual domains to homotypic protein interactions.

### Full-length FoxP1 exhibits increased homotypic interactions than truncated fragments

To quantify FoxP1 homotypic interactions in living cells, we co-transfected HEK293T cells with eGFP- and mCherry-labeled FoxP1 constructs at donor:acceptor (D:A) ratios of 1:1 or 1:5 while maintaining a constant amount of transfected DNA. Cells expressing donor-only (1:0) constructs served as controls. Fluorescence signals from directly excited eGFP and mCherry confirmed successful co-expression of donor- and acceptor-tagged FoxP1 variants and their nuclear localization (**Figure 3A**, left and middle panels). Across all constructs, FoxP1 displayed a heterogeneous intranuclear distribution.

**Figure 3.**
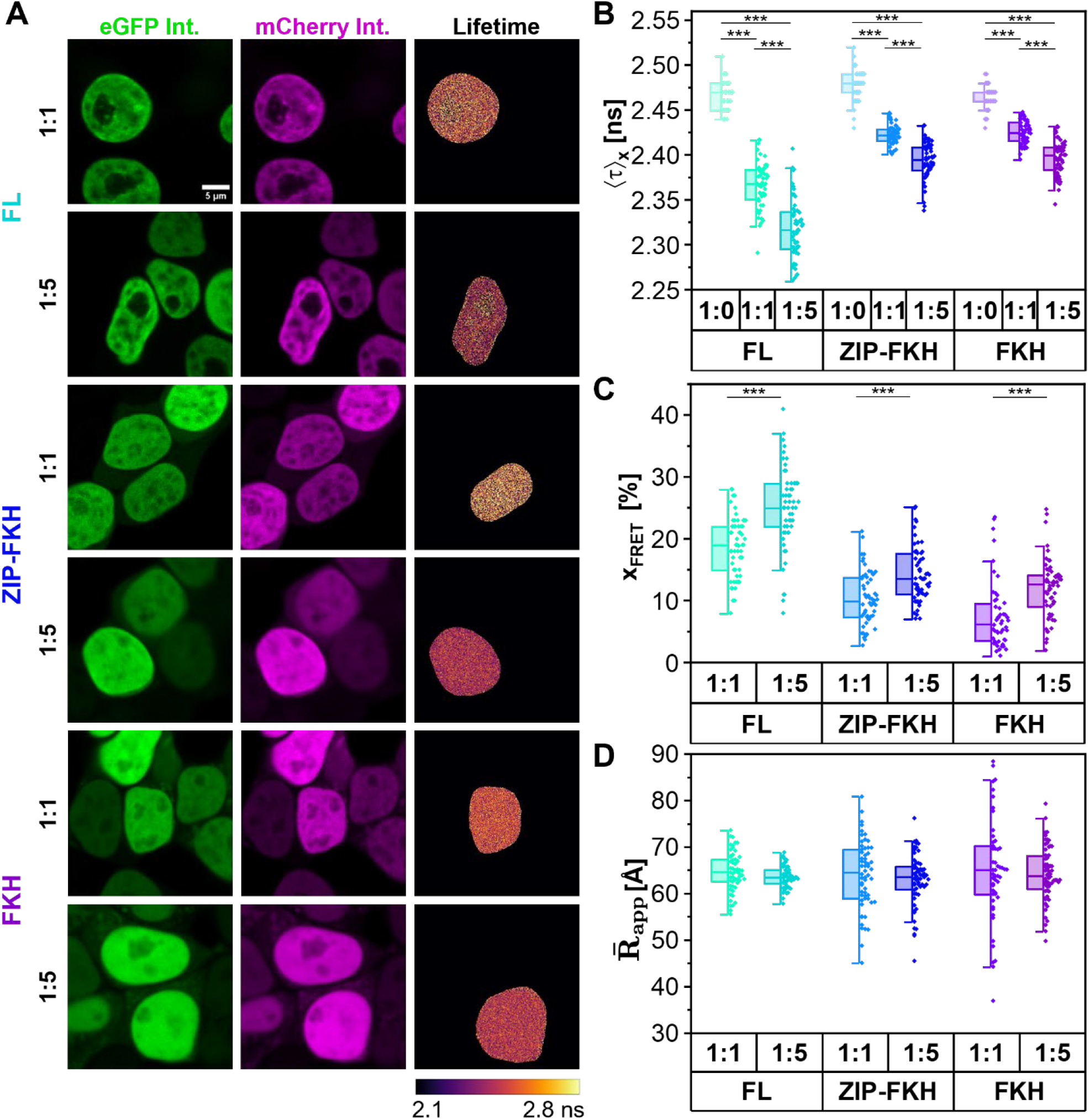
FoxP1 variants exhibit distinct homotypic interaction propensities in HEK293T nuclei. (**A**) Representative PIE-FLIM images of HEK293T cells co-expressing eGFP- and mCherry-tagged FoxP1 constructs at donor:acceptor (D:A) ratios of 1:1 or 1:5. Intensity images shown correspond to: eGFP signal in the prompt time window (left), mCherry signal in the delay time window (middle) and pixel-wise mean arrival time of eGFP photons (right). Image contrast was adjusted for visualization. (**B**) Species-weighted average donor fluorescence lifetime (⟨*τ*⟩_*x*_, **Equation 5**) for the indicated constructs at different D:A ratios (**C**) Fraction of donor molecules undergoing FRET (*x*_*FRET*_) obtained by fitting the donor fluorescence decays with a Gaussian distribution model of donor-acceptor distances (**Equations 6-8**). (**D**) Mean apparent D-A distance (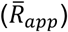 derived from Gaussian distance fitting. Statistical differences between D:A ratios within each FoxP1 variant were assessed using ANOVA.

As an initial assessment of FRET, pixel-wise eGFP fluorescence lifetimes within representative nuclei are shown in **Figure 3A** (right panel). Increasing A:D ratios resulted in a progressive shortening of the eGFP fluorescence lifetime, consistent with enhanced FRET, where increasing the acceptor population raises the probability of forming FRET-active D:A complexes. Notably, donor lifetimes themselves were spatially heterogeneous within individual nuclei, indicating variability in FoxP1 interactions across subnuclear regions. This heterogeneity prompted us to later implement a spatially resolved segmentation strategy to dissect FoxP1 interaction propensities at subnuclear resolution (see below, **Figure 4**)

**Figure 4.**
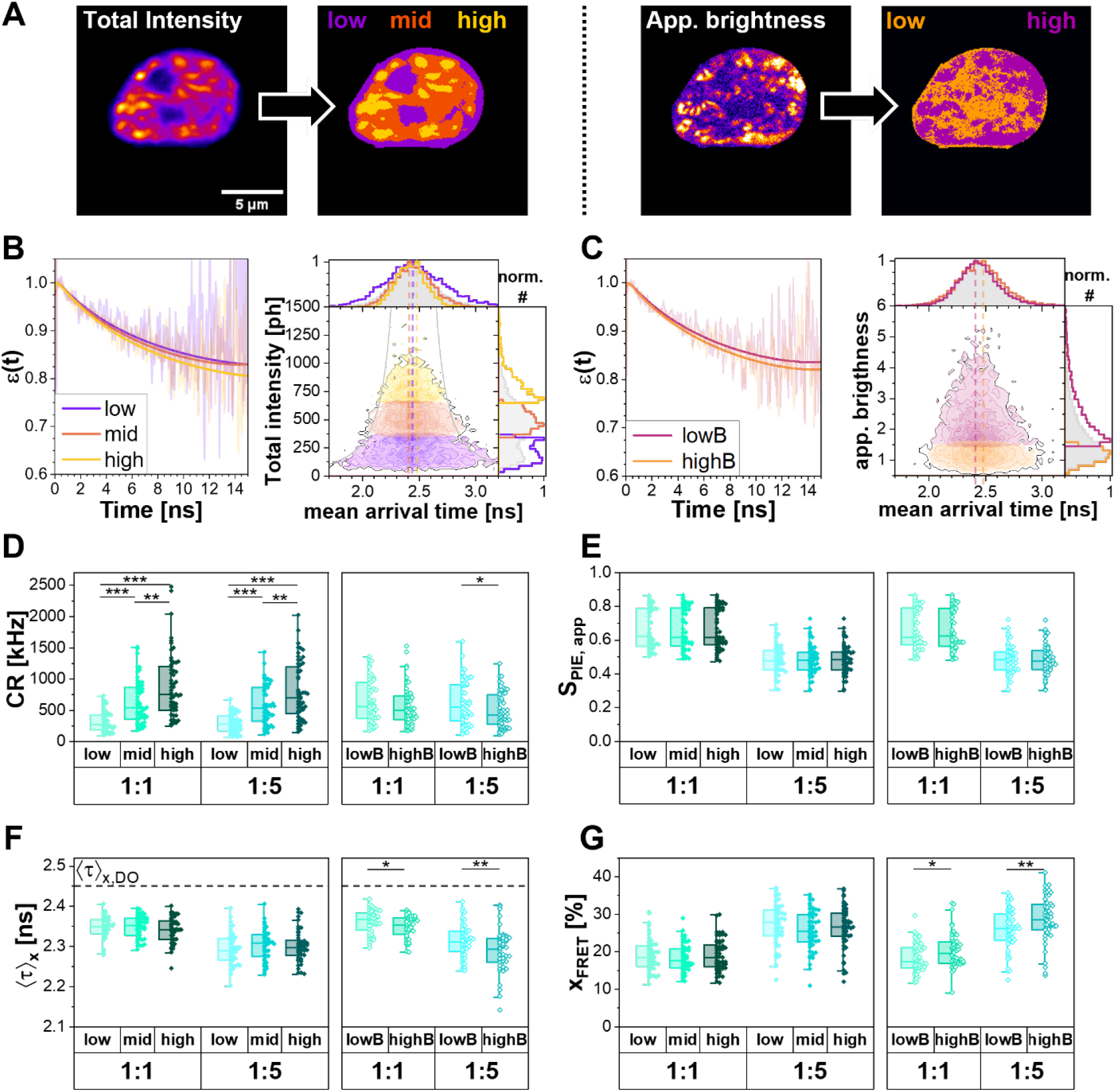
Subnuclear segmentation reveals spatial heterogeneity in FoxP1 homotypic interactions. The FL FoxP1 dataset is shown as representative data. (**A**) Example HEK293T nucleus co-transfected with FL FoxP1 constructs at 1:1 D:A ratio and segmented using the intensity-based (left) or N&B-based (right) approach. (**B**) FRET-induced donor decay, *ε*(*t*), for nuclear regions defined by intensity-based segmentation (solid lines represent fit). Pixel-wise 2D histograms show the three intensity classes (low, mid, high) and their distributions of donor mean arrival time (dashed lines mark the peaks of each distribution) and photon number per pixel. Grey lines mark the expected shot-noised broadening of the mean arrival time distribution at low photon numbers. **C**, Corresponding FRET-induced donor decay and pixel-wise 2D histogram for regions defined by N&B-based segmentation. (**D)** Background-corrected total photon count rate, *CR*, for each subnuclear region. (**E**) Apparent stoichiometry (*S*_*PIE,app*_) for the corresponding regions (**F**) Species-weighted average donor fluorescence lifetime, ⟨*τ*⟩_*x*_ (**Equation 5**). The dashed grey line indicates the average donor-only control lifetime. (**G**) Fraction of molecules undergoing FRET, *x*_*FRET*_, derived from the Gaussian distance fit (**Equations 6-8**). Statistical differences among sub nuclear regions within each D:A ratio and segmentation approach were assessed by ANOVA.

To quantify energy transfer, we defined the nuclei as ROIs and reconstructed time-resolved donor fluorescence decays from the microtime distributions of photons emitted by directly excited donors across all pixels within these regions, as illustrated in **Figure 2B**. These decays were fit using a multiexponential model (**Equation 5**) to determine the species-averaged donor fluorescence lifetime, ⟨*τ*⟩_*x*_, for each construct (**Figure 3B**). All the FoxP1 variants displayed shorter ⟨*τ*⟩_*x*_ at higher acceptor concentrations, reflecting either a larger population of FRET-active complexes, a shorter donor-acceptor (D-A) distance, or a combination of both.

To further distinguish whether these changes arose from altered interaction frequency or structural rearrangements, fluorescence decays were additionally analyzed using a Gaussian distribution model of D-A distances. This analysis yielded an apparent D-A distance, 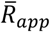, - reporting on the average spatial separation between the C-terminal fluorescent protein tags on interacting FoxP1 molecules, and therefore sensitive to the relative orientation and compactness of FoxP1 complexes – its distribution width, *σ*_*app*_ (**Equation 8**), and the fraction of donor molecules undergoing FRET, *x*_*FRET*_ (**Figure 3C**), a measure of interaction propensity. While *x*_*FRET*_ increased strongly with the acceptor concentration, the average D-A distance (**Figure 3D**) remained largely unchanged. The stability of 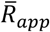 across constructs and conditions, therefore, indicates that the spatial arrangement of FoxP1 in the FRET-active FoxP1 complexes is preserved, consistent with the formation of FoxP1 dimers, and that changes in *x*_*FRET*_ primarily reflect differences in how frequently FoxP1 molecules interact rather than how they are organized when they do. Note that the large spread in 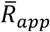 is likely caused by the low *x*_*FRET*_ for the 1:1 ZIP-FKH and FKH samples^59^.

Consistent with that interpretation, FL FoxP1 exhibited the strongest FRET effect, with ⟨*τ*⟩_*x*_ approximately 6% shorter than the donor-only control and *x*_*FRET*_ value roughly twofold higher than that of the truncated fragments at comparable D:A ratios. The ZIP-FKH fragment displayed intermediate ⟨*τ*⟩_*x*_ and *x*_*FRET*_ values, whereas the FKH fragment showed the weakest interaction signature with ⟨*τ*⟩_*x*_ values approaching donor-only controls and an *x*_*FRET*_ of only 7.7%. Together, these data reveal a progressive attenuation of energy transfer efficiency from FL to ZIP–FKH and finally to FKH, consistent with a progressive reduction in homotypic interaction propensity. The enhanced interaction propensity of FL FoxP1 relative to the ZIP-FKH fragment reflects contributions from the ZF domain and Q-rich region, both of which are present in the full-length construct but absent in the truncations. Therefore, these findings indicate that multiple domains cooperate to stabilize FoxP1 homotypic complexes in living cells.

### Subnuclear regions exhibit distinct FoxP1 homotypic interactions

Given the substantial ⟨*τ*⟩_*x*_ and *x*_*FRET*_ variability across constructs (**Figure 3B, C**) and the heterogeneous nuclear FoxP1 distribution (**Figure 3A**), we wondered whether these variations were correlated. To reduce intra-nuclear ensemble averaging and refine FoxP1 interaction analysis, we implemented two image-based segmentation strategies that partition each nucleus into distinct regions (**Figure 4A**): *(i)* intensity-based and *(ii)* the number-and-brightness (N&B)-based segmentation.

In the intensity-based segmentation, each pixel was assigned a compound total fluorescence intensity derived from photons collected across all detectors. This intensity map was segmented using multi-Otsu thresholding^60^, yielding three spatial pixel classes corresponding to low-, mid-, and high-intensity regions (**Figure 4B**).

In parallel, the number-and-brightness (N&B) analysis evaluated total fluorescence intensity fluctuations per pixel as variations in the number of photons collected across all detectors and imaged frames. The apparent brightness (*B)* was calculated as the ratio of variance to mean fluorescence intensity (**Equation 3**), and the apparent number (*N)*, as the ratio of mean fluorescence intensity to *B* (**Equation 2**) ^61^. Monomeric proteins typically exhibit brightness values close to 1, whereas oligomers display higher values. Based on the observed brightness distribution, we applied a fixed threshold of 1.5 to separate the nucleus into low- and high-apparent brightness regions (low-B and high-B, **Figure 4C**). Note, the mean photon arrival time follows a shot-noise distribution^62^; therefore, the broadening at low photon numbers is solely associated with shot-noise.

After defining the five nuclear regions using both segmentation approaches, we calculated the total fluorescence count rate (*CR*) as photon counts per unit time (see data analysis in Materials and Methods) as a proxy for the total FoxP1 concentration, and determined the apparent stoichiometry (*S*_*PIE,app*_), which reports the donor-to-total protein fraction (**Equation 9**) for each region. Both *CR* and *S*_*PIE,app*_ were monitored across segmented nuclear regions to ensure that the segmentation procedure did not introduce artifactual differences in fluorescence measurements. As expected, *CR* increased with intensity, reflecting higher FoxP1 concentration while remaining similar between low- and high-B regions, suggesting no correlation between brightness differences and protein concentration (**Figure 4D**). In both segmentation schemes, stoichiometry was uniform across subnuclear regions and increased with higher acceptor-to-donor ratios (**Figure 4E**), suggesting that regional differences in the measured observables likely reflect biophysical effects rather than differences in donor-acceptor encounter probability.

While ⟨*τ*⟩_*x*_ varied with the D:A ratio, no significant differences were observed among the intensity-based regions. In contrast, high-B regions displayed significantly shorter ⟨*τ*⟩_*x*_ values than low-B regions, independent of the D:A ratio. This reduction indicates greater FRET effect and suggests that the N&B segmentation separates nuclear regions with distinct FoxP1 interaction propensities.

To further characterize the FoxP1 homotypic interaction propensity, we fitted the fluorescence decays by D-A distance distribution models (**Equation 8**). Across the different nuclei, the average D-A distance did not vary (**Figure 3D**). Thus, we fixed the mean distance in the model at 64.2 Å (value derived from the initial Gaussian fit of the FL FoxP1 constructs) and joined the distributions’ widths. Notably, *x*_*FRET*_ remained unchanged across intensity-based regions (**Figure 4G**) but increased significantly in high-B regions relative to low-B regions for both D:A ratios. These results indicate an increased fraction of FoxP1 interacting molecules in high-B regions and demonstrate that N&B-based segmentation partitions the nucleus into distinct FoxP1 homotypic interaction propensities. Moreover, due to the implications of FoxP1 association levels on apparent brightness values (described above), high-B regions present a higher probability of containing not only FRET-active interactions (e.g., dimers or oligomers driven by the C-terminal domains and regions) but also high-order FRET-inactive complexes (e.g., oligomers mediated by the N-terminal), while low-B pixels represent regions with high content of monomers and a fraction of FRET-active dimers.

Together, these findings reveal a spatially heterogeneous FoxP1 self-interaction network within the nucleus.

### FoxP1 oligomerization is mediated by ZIP- and FKH-dependent mechanisms

We applied the N&B-based segmentation to all previously analyzed nuclei expressing the FL, ZIP-FKH and FKH FoxP1 variants to define low- and high-B nuclear subregions. Within each region, the donor fluorescence decays were analyzed using a distance model with fixed 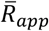 and floating *σ*_*app*_, as described above.

However, the FoxP1 constructs exhibited broad distributions of *S*_*PIE,app*_ and *CR*, spanning 0.2-0.95 and 80-10,000 kHz, respectively (**Supplementary Figure 2**). *x*_*FRET*_, our proxy of homotypic FoxP1 interaction propensity, strongly depends on both parameters (**Figure 5A I, Supplementary Figure 3**, and **Supplementary Table 2**). Thus, for robust cross-construct comparisons, we stratified the nuclei data based on *S*_*PIE,app*_ and *CR* (**Figure 5A, II-IV; details in Methods**). This filtering retained nuclei within comparable ranges of protein abundance and donor-acceptor balance across all constructs.

**Figure 5.**
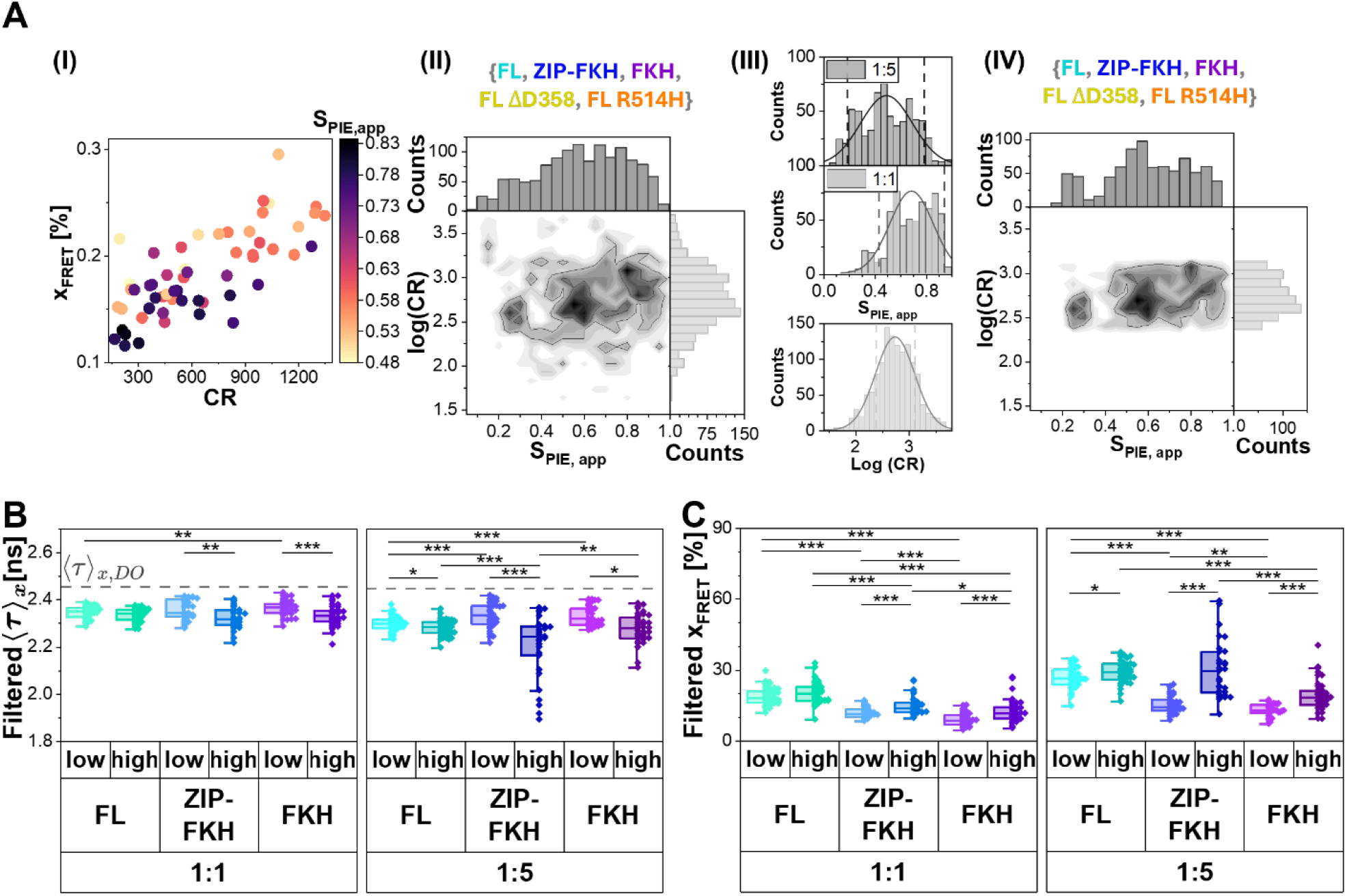
Data stratification reveals distinct self-association behaviors of FoxP1 variants. (**A**) Nuclei data was stratified based on FoxP1 expression parameters. (I) Dependence of *x*_*FRET*_ on apparent stoichiometry, *S*_*PIE,app*_, and count rate, *CR*, illustrated using data from low-brightness regions of cells expressing FL FoxP1 at a 1:1 ratio. (II) 2D histogram of *CR* and *S*_*PIE,app*_ across all five main constructs analyzed in this study (FL, ZIP-FKH, FKH, FL-ΔD358, and FL-R514H). (III) Independent distributions of *CR* and *S*_*PIE,app*_ with Gaussian fits (solid lines) and data-inclusion windows (dashed lines). (IV) 2D histogram of nuclei retained after applying the selection criteria. (**B**) Species-weighted average donor fluorescence lifetime of selected data. Grey dashed line indicates the average mean fluorescence lifetime of the donor-only control groups. (**C**) Fraction of FRET-active FoxP1 determined from Gaussian fitting of D-A distance distributions. Statistical significance was assessed by ANOVA to compare (i) differences among constructs within equivalent brightness regions and (ii) differences between brightness regions within each construct, with analysis performed separately for each D:A ratio.

Filtered ⟨*τ*⟩_*x*_ ranged 2.21-2.43 ns for all constructs at a 1:1 D:A ratio in both brightness regions (**Figure 5B**, left panel), remaining consistently below the donor-only control value. Under these conditions, no significant differences in ⟨*τ*⟩_*x*_ were observed among constructs, except for the FKH fragment, which presented a slightly longer ⟨*τ*⟩_*x*_ when compared to FL FoxP1 in the low-B region.

Consistent with this observation, analysis of *x*_*FRET*_ (**Figure 5C**, left panel) revealed that the FKH fragment displayed the lowest interaction propensity (*x*_*FRET*_∼ 9 % and 13% at low- and high-B, respectively), whereas the FL construct exhibited the highest values (∼ 20% on average). These results agree with whole-nucleus measurements (**Figure 3C**) and indicate stronger homotypic interactions for full-length FoxP1.

At the 1:5 D:A ratio, similar trends were observed (**Figure 5B,C**, right panel). In the low-B region, FL FoxP1 exhibited the shortest ⟨*τ*⟩_*x*_ (∼2.3 ns) and largest *x*_*FRET*_ (27%), confirming a higher interaction propensity relative to the truncated fragments. The ZIP-FKH fragment displayed complex behavior: while lifetime remained relatively long in the low-B region (∼2.33 ns), the high-B region exhibited shorter lifetimes (∼2.20 ns) with greater dispersion. Consequently, the mean *x*_*FRET*_ of ZIP-FKH and FL FoxP1 were similar in high-B regions.

Across all conditions, the FKH fragment consistently displayed the lowest non-zero *x*_*FRET*_, in agreement with its weak dimerization propensity observed *in vitro* (**Figure 1D**). Collectively, these results reveal a markedly higher frequency of ZIP-FKH homotypic interactions compared with those driven by the FKH domain alone. Consistent with previous studies on isolated protein^28^, these findings indicate the presence of two distinct FoxP1 self-association mechanisms in living cells.

### Disruption of DNA binding enhances FoxP1 homotypic interactions

To investigate how FoxP1 homotypic interactions are affected by pathogenic structural alterations, we introduced point mutations into the FL FoxP1 sequence. The R514H mutation is a missense variant within the FKH domain that disrupts FoxP1-DNA binding^36,38^. The FL-ΔD358 mutant involves the deletion of aspartic acid 358 within the leucine zipper domain. Deletion of this residue is predicted to disrupt the heptad-repeat periodicity of the coiled-coil interface, misaligning the leucine residues that mediate ZIP-dependent dimerization and thereby destabilizing this interaction, as observed for a similar mutation^32^. This mutation was therefore selected as a tool to specifically interrogate the contribution of ZIP-mediated dimerization to the full-length FoxP1 complex.

HEK293T cells were transfected with the mutant constructs, and their nuclear localization was examined (**Supplementary Figure 1**). While FL-ΔD358 displayed clear nuclear localization, the FL-R514H showed a punctate distribution either outside or within the nucleus. For this mutant, nuclei were manually delineated prior to analysis. All datasets were then processed using the established workflow, including N&B-based segmentation (**Figure 6A**) and filtering based on *S*_*PIE,app*_ and *CR* to ensure comparability across samples.

**Figure 6.**
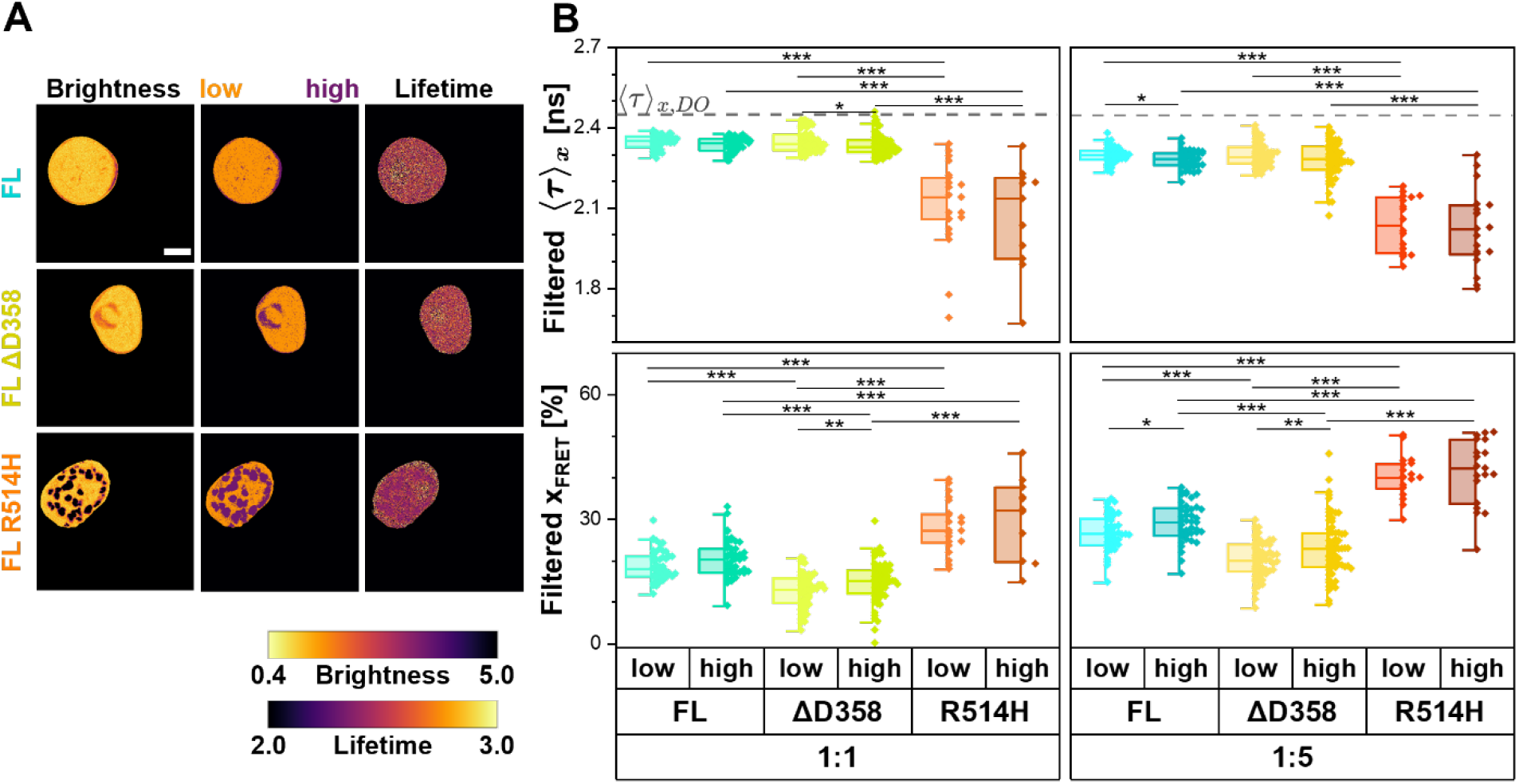
Pathogenic FoxP1 mutations differentially affect homotypic interactions. (**A**) Representative nuclei expressing FL, FL ΔD358 and FL R514H constructs (1:1 ratio). Images show apparent brightness (left), subnuclear regions defined by N&B-based ROI segmentation (middle), and pixel-wise donor fluorescence lifetimes (right). The scale bar represents 5 μm. (**B**) Species-weighted donor fluorescence lifetime ⟨*τ*⟩_*x*_ (top) and fraction of FRET-active molecules *x*_*FRET*_ (bottom) for FL, FL ΔD358 and FL R514H variants after filtering based on *S*_*PIE,app*_ and *CR* criteria defined in **Figure 5**. The grey dashed line indicates the average donor-only lifetime. Statistical significance was assessed by ANOVA to compare (i) differences among constructs within equivalent brightness regions and (ii) differences between brightness regions within each construct. All analyses were performed separately for each D:A ratio.

The FL-ΔD358 mutant exhibited reduced *x*_*FRET*_ values (**Figure 6B**, bottom), indicating a decreased homotypic interaction propensity. This observation is consistent with previous *in vitro* studies demonstrating that the ZIP domain is essential for FoxP1 dimerization^32^. However, the ΔD358 mutation did not completely abolish FoxP1 interactions, as ⟨*τ*⟩_*x*_ remained below the donor-only control value, and *x*_*FRET*_ exceeds 10%. These residual interactions suggest that the ZIP domain retained partial functionality or that additional regions contributed to the formation of the full-length FoxP1 complex, consistent with our earlier observations (**Figure 5**).

In contrast, the FL-R514H mutant exhibited the shortest ⟨*τ*⟩_*x*_ (∼ 2.06 ns on average) and the highest *x*_*FRET*_ (∼ 29% and 41% for 1:1 and 1:5 D:A ratios, respectively) across both D:A conditions and nuclear regions (**Figure 6B**), strongly indicating an enhanced FoxP1 homotypic interaction propensity which correlates with a decreased DNA-binding affinity of the FKH domain.

This increased interaction propensity is consistent with the punctate fluorescence pattern seen in cells transfected with FoxP1 R514H (**Figure 6A**, bottom), a feature commonly associated with biomolecular condensates^63^. Notably, despite the presence of an NLS in all FoxP1 constructs, this mutant frequently failed to accumulate efficiently in the nucleus (**Supplementary Figure 4**).

Analysis of the time-resolved fluorescence decays revealed that the R514H mutant required fitting with two Gaussian distance components (**Equation 8**). The longer D-A distance (*R*_*app,1*_ = 66.1 Å) was comparable to that observed in all other constructs, consistent with dimeric FoxP1 complexes. The shorter distance (*R*_*app,2*_ = 33.4 Å) is consistent with multi-acceptor FRET arising from dense molecular packing within the observed puncta, where the proximity of multiple acceptor molecules to a single donor produces an increased apparent FRET effect and a correspondingly shortened apparent distance^64^. This interpretation is supported by the punctate fluorescence distribution of R514H (**Figure 6A**) and is consistent with previously reported multi-acceptor FRET signatures in densely packed protein assemblies^65^. Accordingly, the total *x*_*FRET*_ value was defined as the sum of the contributions from both distance components.

### FoxP1 oligomers accumulate at the nuclear periphery

Given the abnormal cellular distribution of the FL-R514H mutant, we investigated whether the nuclear localization of FoxP1 complexes depends on underlying interaction mechanisms. To this end, we defined a peripheral nuclear rim of 0.5 μm width for all nuclei (**Figure 7A**), corresponding on average to 19% of the total nuclear area. This boundary was based on the normalized radial density profiles of high brightness pixels across all constructs (**Figure 7B**, and **Supplementary Figure 5A-C**). Owing to its aberrant localization, the FL-R514H mutant was excluded from this analysis.

**Figure 7.**
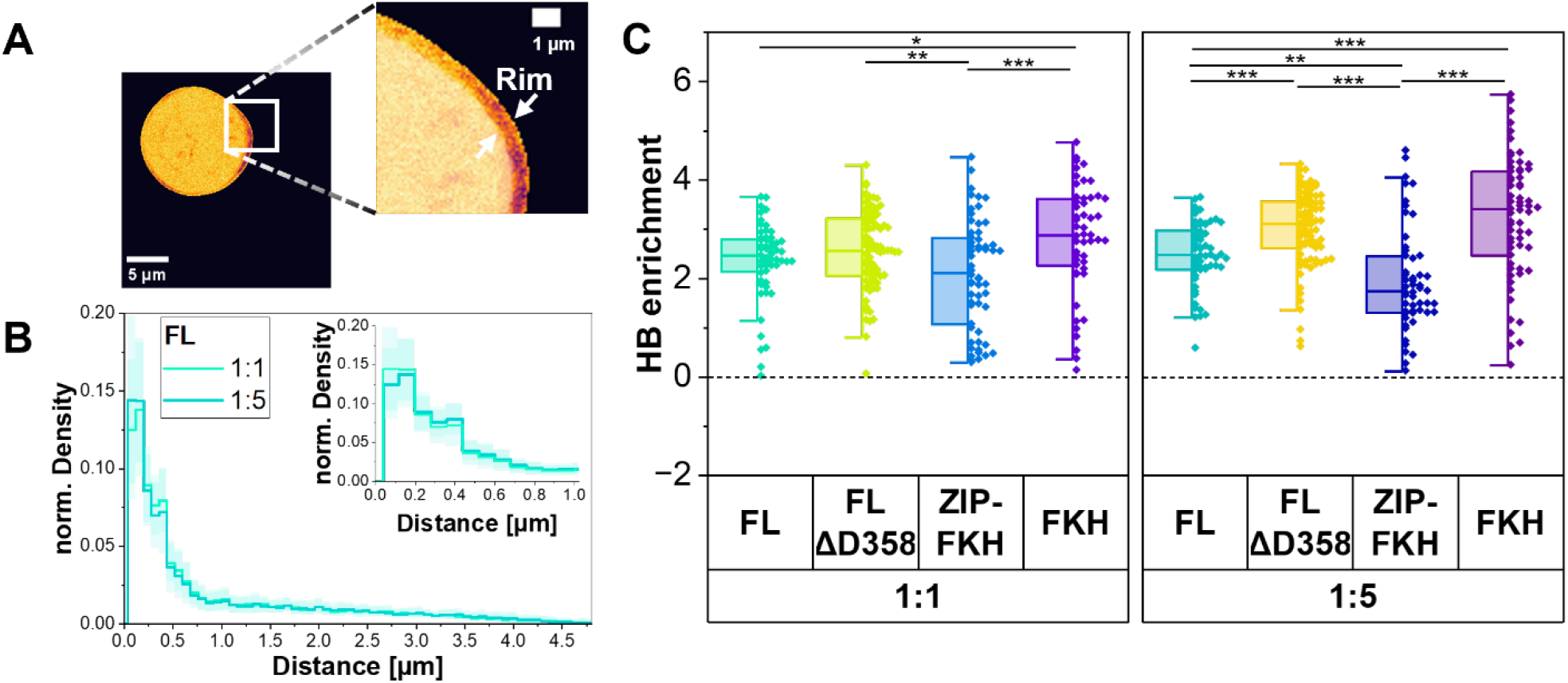
FoxP1 oligomers preferentially accumulate at the nuclear periphery. (**A**) Representative apparent brightness image illustrating the definition of the peripheral rim (0.5 μm width). (**B**) Normalized radial density of high-brightness pixels relative to the nuclear border (0 µm) for FL FoxP1. (**C**) Enrichment of high-brightness (HB) pixels in the rim relative to the nuclear center. The black dashed line indicates equal distribution. Values > 0 indicate enrichment in the nuclear rim.

Across constructs, FL FoxP1 exhibited the largest fraction of high-brightness pixels within the nucleus (∼16% in average) (**Supplementary Figure 5D**). In contrast, the other variants showed reduced fractions of high-B pixels, including FL-ΔD358 (∼11%), FKH (∼8%), and ZIP-FKH (∼3%). A similar trend was observed within the nuclear rim: FL FoxP1 showed the highest proportion of high-brightness pixels (∼ 50% in average), whereas FL-ΔD358 (∼40%) and FKH (∼33%) showed moderate reductions, and ZIP-FKH (∼12%) exhibited a pronounced decrease (**Supplementary Figure 5E**).

To dissect the spatial organization of distinct FoxP1 species, we defined the high-B pixel-enrichment (**Figure 7C**) as the logarithm of the ratio between the fraction of high-B pixels in the rim and the fraction of high-B pixels in the nuclear center (**Equation 10**). Considering that high-B pixels contain brightness-defined oligomers and other higher-order configurations (e.g., loosely packed condensates that may fall below the FRET detection threshold due to large D-A distances) and just a small fraction of FRET-active dimers, and that *x*_*FRET*_ showed only marginal differences between distinct brightness regions for most constructs and experimental conditions, this parameter captures the spatial distribution of higher-order FoxP1 complexes and allows us to spatially decouple them from dimeric FoxP1. Therefore, positive values of HB enrichment indicate enhanced concentration of higher-order FoxP1 complexes in the rim, zero indicates uniform distribution across the nucleus, and negative values indicate enrichment in the nuclear center.

Across constructs, high-B pixels were strongly enriched at the nuclear periphery (**Figure 7C**), with positive log_2_ values corresponding to several-fold accumulation relative to the center. Furthermore, FKH-mediated complexes accumulate more at the nuclear periphery than ZIP-mediated complexes, with a combinatorial effect on FL FoxP1, suggesting that peripheral FoxP1 complexes are governed by FKH-mediated interactions. Altering ZIP-mediated interactions shifts FL FoxP1 to the characteristic behavior of FKH-mediated complexes, as observed by the similar peripheral enrichment of ΔD358 FL and FKH complexes, suggesting the emergence of compensatory FKH-mediated interactions upon disruption of ZIP-driven complexes.

Collectively, these results reveal a spatial decoupling of FoxP1 complex configurations within the nucleus; with higher-order complexes, preferentially driven by the FKH domain, partitioning to the nuclear periphery; while FRET-active dimers, preferentially driven by the ZIP domain, remain more towards the center of the nucleus. Considering the nuclear differential organization of chromatin structure, FoxP1 spatial distribution suggests that peripheral accumulation reflects its sequestration within higher-order complexes rather than enhanced dimerization activity, and that functionally engaged FoxP1 predominantly operates in a dimeric configuration at central nuclear regions.

## DISCUSSION

In this study, we show that FoxP1 assembles into structurally heterogeneous complexes in living cells through coupling between its folded (ZIP and FKH) and intrinsically disordered domains (Q-rich). Using PIE-FLIM, we demonstrate that FoxP1 does not adopt a single stable configuration but instead samples multiple states whose stability and prevalence depend on domain composition and sequence integrity. This dynamic behavior contrasts with classical models of rigid transcription factor dimers, such as the stable bZIP homodimers of GCN4^29^ or the fixed geometry of p53 tetramers^66^, in which a single dominant interaction interface defines the entire functional complex. Such structural plasticity likely enables FoxP1 to adapt to diverse chromatin environments and regulatory contexts.

PIE-FLIM measurements on truncation variants and previous observations^28^ reveal that FoxP1 homo-complexes are governed by negative interdomain coupling between the ZIP and FKH domains (**Figure 5**). While the ZIP domain promotes dimer formation, it simultaneously suppresses FKH-mediated interactions. This balance enables FoxP1 to tune its interaction state in response to structural constraints and cellular context. A clear example is the disruption of ZIP-mediated interactions in the ΔD358 FL variant, which shifts FL FoxP1 behavior toward a FKH-mediated complex (**Figure 7C**).

Notably, comparison of *in vitro* dimerization assays (**Figure 1**) with live-cell FRET measurements (**Figure 3**) highlights the importance of additional cellular and molecular factors that facilitate FoxP1 homotypic interactions. The *in situ* enhanced interaction propensity of FL FoxP1 relative to the ZIP-FKH fragment suggests contributions beyond the ZIP and FKH domains. The Q-rich, a low-complexity region, is a prime candidate, given its proposed secondary structure (**Figure 1**) and the established role of such domains in promoting protein-protein contacts and condensate formation in other transcription factors^53,54^. However, its specific contribution to FoxP1 assembly requires direct experimental investigation and represents an important direction for future work

Multiple FoxP1 complex configurations have significant consequences for DNA binding and chromatin interactions. Previous structural studies proposed that FKH domain swapping might enable chromatin bridging and repression through stable dimer formation^36^. Several observations in our study suggest flexible, spatially organized, diverse binding modes. While brightness-defined FoxP1 oligomers preferentially accumulate at the nuclear periphery (**Figure 7C**), the fraction of FRET-active molecules shows only minor variation across the nucleus (**Figure 5C** and **6B**). Given that FRET-active species likely report on dimers, this distribution suggests that oligomeric FoxP1 (or another higher-order configuration) interacts with heterochromatin, the dormant chromatin structure that preferentially localizes at the nuclear periphery. This potential interaction supports FoxP1’s role as a pioneer TF, enabling it to access compacted chromatin and reorganize local regulatory landscapes. Consistently, this potential behavior implies that FoxP1 dimers regulate gene expression through a more traditional mechanism by interacting with euchromatin. Together, these implications support a model in which FoxP1 engages chromatin through transient, spatially regulated self-interactions rather than a single rigid architecture.

Critically, neither the disease-associated R514H nor the ΔD358 deletion variant acts by simply abolishing a single interaction interface; instead, both perturb the interdomain balance that governs FoxP1’s assembly spectrum, shifting the equilibrium toward either hyperstabilized condensates or reduced dimerization (**Figure 6**). This distinction has broader implications: it suggests that pathogenic variants in multidomain transcription factors may commonly act by disrupting interdomain coupling rather than by loss of a single functional domain, a mechanism that would be missed by approaches that examine individual domains in isolation. These findings provide a mechanistic link between sequence alterations, defective assembly, and transcriptional dysregulation. More broadly, they illustrate a model of how subtle changes in domain coupling can have profound functional consequences in multidomain transcription factors.

A key enabler of these findings was the combination of PIE-FLIM with number-and-brightness analysis (**Figure 4**), which allowed us to resolve spatially heterogeneous FoxP1 interaction states within intact living cell nuclei - a distinction that would be inaccessible to ensemble biochemical approaches. Specifically, the ability to simultaneously quantify interaction frequency, molecular brightness, and spatial distribution within the same nucleus revealed that FoxP1 complexes are not uniformly distributed but are spatially organized in a domain-dependent manner. The sensitivity of this approach to weak and transient interactions, combined with its robustness to variable expression levels and spectral cross-talk, makes it broadly applicable to other multidomain nuclear proteins whose organization dynamics have resisted characterization by conventional methods.

FoxP1 exemplifies a growing class of transcription factors whose regulatory activity emerges from the interplay of multiple folded domains and intrinsically disordered regions. Our findings suggest that the encoding of multiple functional complexes within a single polypeptide - governed by competing interaction interfaces rather than a single dominant dimerization mode - represents a fundamental and likely widespread design principle of transcriptional control. Consistent with this view, related Fox family members, including FOXP2 and FOXP4, share the ZIP and FKH domain architecture implicated here^32^, suggesting that interdomain coupling may similarly regulate their assembly and function. More broadly, the principle that multidomain architecture encodes a spectrum of regulatory states, rather than a single functional conformation, may apply across transcription factor families that combine structured oligomerization domains with intrinsically disordered regions - a combination that characterizes the majority of eukaryotic transcriptional regulators^16^.

Together, our results demonstrate that FoxP1 operates through multiple heterogeneous complexes whose organization is governed by interdomain coupling. We further show that these complexes are spatially organized within the nucleus, with oligomers enriched at the nuclear periphery, whereas FRET-active dimers remain more uniformly distributed. This spatial decoupling suggests that higher-order complexes and functional DNA engagement are distinct yet coordinated features of FoxP1 gene regulation. By revealing how structural flexibility and modular interactions shape FoxP1 function in living cells, this work advances our understanding of transcription factor regulation and provides a foundation for investigating how dysregulated complexes contribute to human disease.

### Limitations of the study

This study establishes a quantitative framework for analyzing FoxP1 assembly in living cells while highlighting directions for future work. First, our experiments used fluorescently tagged, overexpressed constructs in HEK293T cells, which do not endogenously express high levels of FoxP1 and may not fully recapitulate the chromatin and cofactor environment of physiologically relevant cell types such as B cells, neurons, and cardiac progenitors. Extension to endogenously tagged FoxP1 in native cellular contexts will be important for validating these findings. Second, our measurements focus on FoxP1 homotypic interactions and do not directly report chromatin state or cofactor interactions. Incorporation of orthogonal chromatin reporters and multicolor imaging approaches will enable future studies to link FoxP1 assembly states to specific chromatin environments and cofactor recruitment. Third, as with all FRET-based methods, our analysis is sensitive to molecular proximities within approximately 7–8 nm, and distances are reported as apparent values assuming κ^2^ = 2/3 and R_0_ = 52 Å. While these assumptions are standard and the comparative conclusions of this study are robust to small variations in these parameters, combining PIE-FLIM with complementary structural techniques will further refine our understanding of FoxP1 architecture. Additionally, the truncated constructs used for live-cell measurements retain the C-terminal region absent in the corresponding *in vitro* preparations, meaning absolute interaction parameters are not directly comparable across experimental systems, though the relative ordering of interaction propensities remains consistent. Finally, linking specific assembly states to genome-wide binding patterns and transcriptional outputs will require combining PIE-FLIM with chromatin profiling and functional assays.

## RESOURCE AVAILABILITY

### Lead contact

Further information and requests for resources and reagents should be directed to and will be fulfilled by the lead contact, Katherina Hemmen.

### Materials availability

Plasmids newly generated in this work are available upon request to the lead contact. No unique reagents were generated for this study.

### Data and code availability

Datasets and scripts are available on Zenodo (https://doi.org/10.5281/zenodo.18347876, https://doi.org/10.5281/zenodo.18360348, https://doi.org/10.5281/zenodo.18362153). Step-by-step protocols for the software and scripts used have been deposited within the framework of a different project (https://doi.org/10.5281/zenodo.17869433).

## ACKNOWLEDGMENTS

The microscopy was supported by the Interdisciplinary Center of Clinical Research (IZKF) of the Medical Faculty of Würzburg [Grant No. Z-12 to KGH] and by the DFG (Deutsche Forschungsgemeinschaft) by funding the LSM980 (DFG-INST 93/1022-1).

The authors acknowledge support from NIH [Grant No. CA2080699 and GM151334], Clemson University, Mexican Secretaría de Ciencia, Humanidades, Tecnología e Innovación (SECIHTI) through the program “Becas de Posgrado en Ciencia y Humanidades en el Extranjero 2025”, and Chilean National Agency of Investigation Development (FONDECYT 1251879, FONDEQUIP EQM200202).

We thank Kerstin Jansen for help with cell culture and Mike Friedrich for technical support on the LSM980.

## AUTHOR CONTRIBUTIONS

A.L. and K.H. created the mammalian plasmids and performed the PIE-FLIM experiments. T.O.P., K.H. and H.S. developed the PIE-FLIM analysis workflow. T.O.P. and K.H. wrote analysis software and scripts. A.L. and K.H. analyzed the PIE-FLIM data. E. M. and J.A performed and analyzed the in vitro experiments. E.M., K.H., K.G.H., and H.S. devised the study. A.L., K.H. and H.S. wrote the manuscript. All authors contributed to the editing of the manuscript.

## DECLARATION OF GENERATIVE AI AND AI-ASSISTED TECHNOLOGIES IN THE WRITING PROCESS

During the preparation of this work, the authors used ChatGPT and Grammarly in order to corroborate the grammar and coherence of the text. After using this tool, the authors reviewed and edited the content as needed and take full responsibility for the content of the publication.

## SUPPLEMENTAL INFORMATION

Document S1. Figures S1-S6 and Tables S1-S4.

## STAR★METHODS

### KEY RESOURCES TABLE

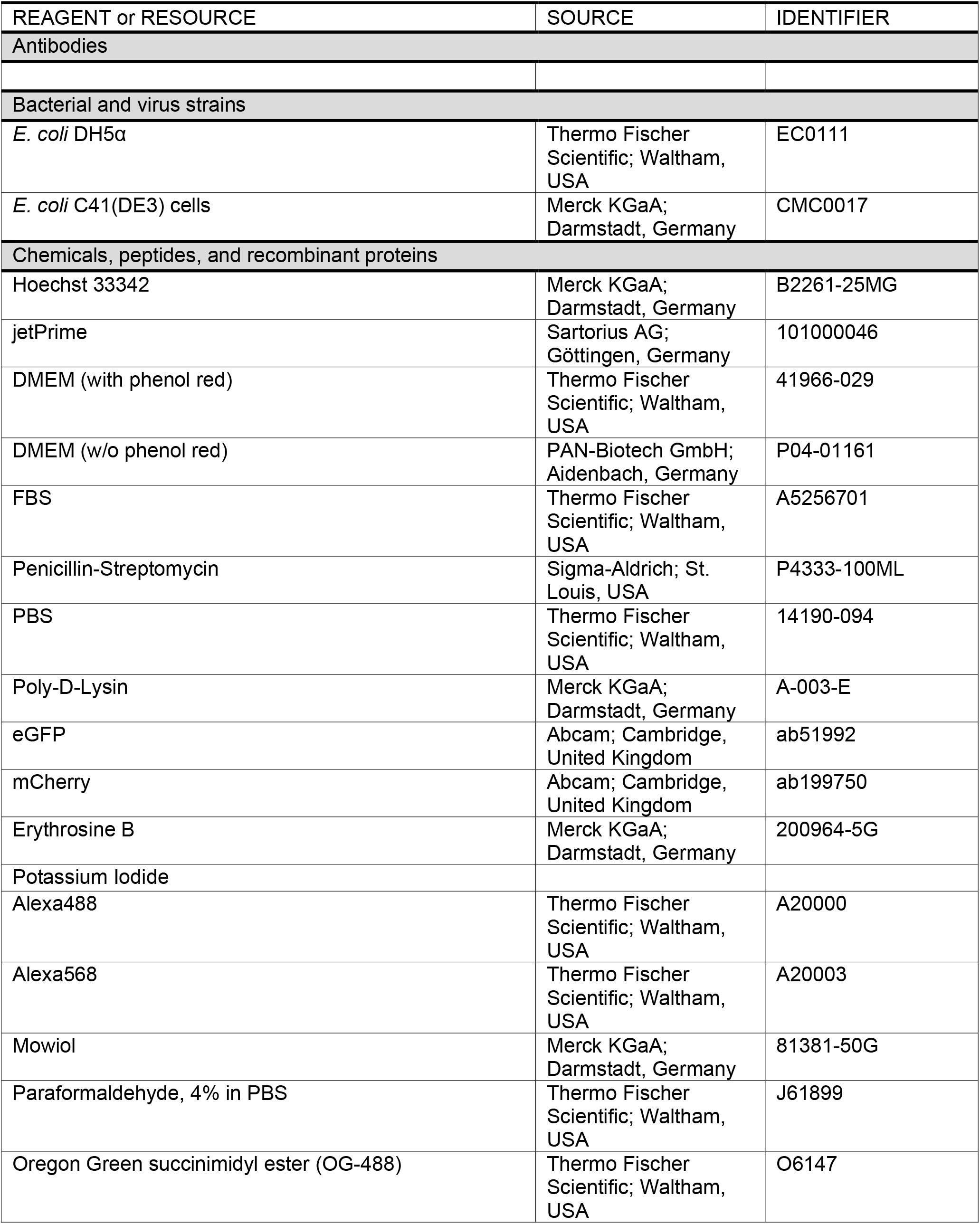
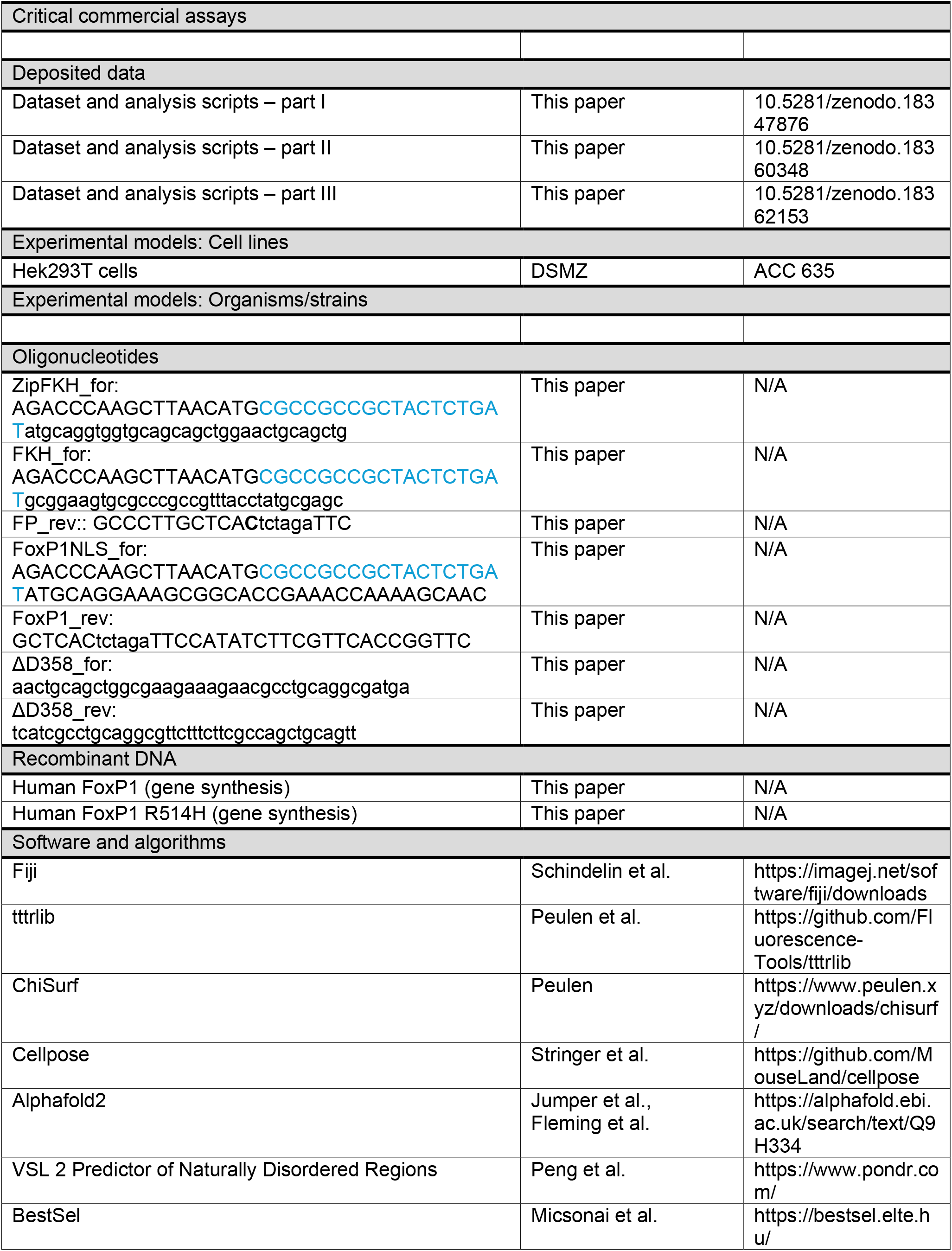
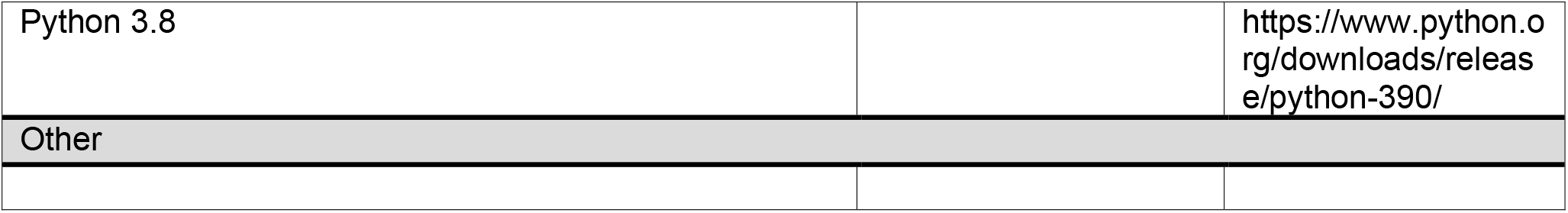

## METHOD DETAILS

### Sample preparation

#### Recombinant FL-FoxP1 purification and labeling

Human full-length FoxP1 (FL-FoxP1) was cloned into a modified pET-28a vector containing an N-terminal His6-tag followed by a TEV protease cleavage site. The construct was transformed into *E. coli* C41 cells (Invitrogen) for protein production. Protein expression was induced with 0.5 mM IPTG, followed by overnight incubation at 25 °C until an optical density at 600 nm (*A600*) of 0.8-0.9 was reached. FL-FoxP1 was purified using Ni^2+^-NTA affinity chromatography, as previously described ^28^.

For fluorescent labeling, purified protein (100 µM) was incubated in buffer A (20 mM HEPES, pH 7.5, 150 mM NaCl, 2 M GdmCl) for 30 min at room temperature. Proteins were labeled with Oregon-Green (OG-488), a succinimidyl-ester reactive dye (Thermo Fisher Scientific, Waltham, USA) at a five-fold molar excess, incubated for 2 h at room temperature, followed by 16 h at 4°C. Unreacted dye was removed by size exclusion chromatography.

#### Plasmid preparation for live-cell experiments

Human wildtype (WT) FL FoxP1 and the FL R514H mutant were synthesized (GeneArt, Life Technologies GmbH, Darmstadt, Germany) and cloned into pcDNA3 between the *HindIII* and *XbaI* restriction sites. Each construct was fused N-terminally to eGFP (donor) or mCherry (acceptor) and contained the FoxP1-derived nuclear localization sequence (RRRYSD). Two truncation constructs were generated: a ZIP-FKH fragment (residues 345-677) and a FKH fragment (residues 462-677). The R514H point mutation and the ΔD358 deletion were introduced into these fragments by overlap extension PCR. All constructs were verified by Sanger sequencing. Plasmid maps and sequences are provided in **Supplementary Figure 6, Supplementary Table 3**-**4**.

#### Cell culture

Human embryonic kidney 293T (HEK293T) cells were cultured in Dulbecco’s modified Eagle medium (DMEM) without sodium pyruvate (P04-03550, PAN Biotech, Aidenbach, Germany), supplemented with 10% fetal calf serum (FCS) (AC-SM-0190, Anprotec, Bruckberg, Germany), penicillin (100 U/mL) and streptomycin (100 µg/mL) (P4333, Sigma-Aldrich, St. Louis, Missouri, United States). Cells were maintained at 37°C and 5% CO_2_ and passaged every 3-4 days. For passaging, cells were washed with PBS (14190, Gibco, Thermo Fisher Scientific, Waltham, Massachusetts, United States) and detached using 0.5× Trypsin-EDTA (T4299, Sigma-Aldrich).

#### Cell transfection

For confocal microscopy, 75’000 HEK293T cells were seeded onto 18 mm coverslips (0111640, Marienfeld-Superior, Lauda-Königshofen, Germany) placed in poly-D-Lysine-coated 12-well plates (140675, Thermo Fisher Scientific, Roskilde, Denmark) 24 h prior to transfection. For PIE-FLIM experiments, the same number of cells were seeded on poly-D-lysine-coated 4-chamber cover glass slides (C4-1.5P, Cellvis, Mountain View, California, United States) 48 h prior to transfection, allowing cells to reach optimal confluency for live imaging, given the lower DNA amounts used in these experiments.

Transient transfections were performed using jetPRIME reagent according to the manufacturer’s instructions (Sartorius Sartorius GmbH & Co. KG, Goettingen, Germany). For confocal microscopy, 0.8 µg total DNA was used per sample, whereas 0.5 µg total DNA was used for PIE-FLIM experiments. Cells were transfected with either donor-tagged construct (donor-only controls) or donor and acceptor constructs combined at donor:acceptor (D:A) ratios of 1:1 or 1:5 (FRET conditions). For PIE-FLIM experiments, the amount of tagged construct did not exceed 0.25 µg to limit overexpression. The total DNA amount was adjusted to 0.5 µg using the empty pcDNA3 vector.

Six to eight hours after transfection, the culture medium was replaced with fresh DMEM, and the cells were incubated overnight before imaging.

### Data acquisition

#### Circular Dichroism (CD) measurements

Far-UV CD measurements were performed using a Jasco J-1500 spectropolarimeter with a 1 mm path-length cuvette. FL FoxP1 was dialyzed against CD buffer (20 mM sodium phosphate, pH 7.8, 0.5 mM TCEP and 50 mM sodium sulfate). Spectra were acquired at a protein concentration of 10 µM by averaging three scans recorded between 200 and 250 nm (scan rate of 100 nm⋅min^™1^). Spectra were smoothed using a Savitzky-Golay filter (10 points, polynomial order 2) in OriginPro 2025b. Secondary structure content was estimated using BestSel^51,52^.

#### *In vitro* association measurements

Protein-protein and DNA-protein association were monitored by steady-state fluorescence anisotropy using a Zeiss LSM710 confocal microscope equipped with a diode laser at 485 nm (LDH-P 485 PicoQuant, Germany; power at objective: 120 μW). Fluorescence from freely diffusing molecules was collected through a 70 μm pinhole using a 40 × 1.2 NA collar-corrected Zeiss objective (correction 0.17). Emission passed through a bandpass (HQ 520/35, Semrock), and was separated into parallel and perpendicular polarization channels. For DNA-protein association, the DNA sequence reported in Cruz *et al*.^28^ was used and labeled with Oregon Green 488 (OG-488).

Association experiments were performed with 20 nM of labeled molecules (FL FoxP1-OG488 or DNA-OG488) in standard buffer (20 mM HEPES, pH 7.8, 20 mM NaCl, 2 mM β-mercaptoethanol). For DNA-protein experiments, labeled DNA was mixed with 180 nM of unlabeled DNA. Protein-protein association was measured by titrating unlabeled FL FoxP1 concentrations (0-30 µM) into FL-FoxP1-OG488. For DNA-protein binding assays, unlabeled FL FoxP1 was titrated from 0 to 5 µM. Samples were incubated for 30 min at 37 °C prior to measurement. The G-factor was determined from measurements of Rhodamine 110.

#### Confocal Microscopy Localization Assay

Twenty-four hours after transfection, cells were washed with PBS, fixed with 4% paraformaldehyde (PFA) (158127, Sigma-Aldrich) for 15 min, and washed again with PBS. Nuclei were counterstained with Hoechst dye (0.33 µg/mL) for 5 min and washed with PBS. Coverslips were mounted on microscope slides using 20 µL Mowiol 4-88 (0713.1, Roth, Karlsruhe, Germany). Images were acquired on a Zeiss LSM980 confocal microscope (Zeiss, Oberkochen, Germany) using a 63×/1.4 NA oil-immersion objective, ZEN3.3 software, voxel dimensions of 35×35×130 nm, and a pinhole diameter of 48 µm (1 Airy Unit) to satisfy the Nyquist criterion. Acquisition settings included bidirectional scanning, 2x line averaging, and a pixel dwell time of 1.4 µs-5.5 µs at 16-bit image depth.

The fluorescence signal was recorded in sequential tracks. eGFP was excited at 488 nm and detected between 499-587 nm using a GaAsP-PMT. In the second track, Hoechst (excitation: 353 nm; emission: 408-495 nm, Multialkali-PMT) and mCherry (excitation: 587 nm; emission: 596-693 nm; GaAsP-PMT) were recorded simultaneously. Laser power and detector gain were adjusted for each sample to avoid detector saturation.

#### Pulsed Interleaved Excitation Fluorescence Lifetime Imaging Microscopy (PIE-FLIM)

Twenty-four hours after transfection, the culture medium was replaced with imaging medium consisting of phenol red-free DMEM (P04-01161, PAN Biotech) supplemented with 15 mM HEPES (15630-080, Gibco), 10% fetal calf serum, and 2 mM glutamine (G7513, Sigma-Aldrich). This medium allows live cell imaging for 2-3 h in the absence of a CO_2_ control.

PIE-FLIM measurements were performed on a Zeiss LSM980 confocal microscope (Zeiss, Oberkochen, Germany) using ZEN3.3 software. The system was equipped with a PicoQuant LSM upgrade kit (PicoQuant, Berlin, Germany) and controlled using SymPhoTime64 software. Confocal imaging and time-resolved fluorescence detection were integrated using the Zeiss Blue PicoQuant Application. Live-cell samples were imaged using a 40×/1.2 NA water immersion objective. Excitation was provided by pulsed diode lasers at 485 nm and 560 nm (LDH-P-C-485B/LDH-D-TA-560B, Picoquant, Berlin, Germany). Emitted photons were first separated by a polarizing beam splitter and then by a 560 nm long pass (560/LPXR, AHF, Tübingen, Germany) to distinguish green and red signals. Green fluorescence was filtered using HC520/35 bandpass filters (Semrock, New York, U.S.A.) and detected with PMA Hybrid 40 detectors (Picoquant, Berlin, Germany). Red fluorescence passed through an ET600/50 bandpass filter (Chroma Technology GmbH, Olching, Germany) and was detected by Excelitas SPAD detectors (SPCM-AQRH-14-TR, Picoquant, Berlin, Germany). Images were acquired at a zoom factor of 5.0 (pixel size: 0.083 µm) with a 512 × 512-pixel resolution. Data was collected with a pixel dwell time of 8.16 µs over 40 frames. The repetition rate was set to 40 MHz. PIE^56^ was implemented with a 25 ns delay and a 50 ns time window to alternate direct donor and acceptor excitation. Excitation laser powers were adjusted between 100-500 nW at 485 nm and 200-600 nW at 560 nm. Time-correlated single photon counting (TCSPC) resolution was 10 ps.

Prior to live-cell imaging, the setup was calibrated using standard procedures. The instrument response function (IRF) was measured using a saturated solution of Erythrosine B (200964, Sigma-Aldrich) in potassium iodide (KI) (221945, Sigma-Aldrich). Daily fluorescence correlation spectroscopy (FCS) and TCSPC measurements of Alexa488, Alexa568, eGFP, and mCherry were used as molecular brightness and diffusion standards to determine the instrumental G-factor and polarization mixing factors^67,68^, and to correct for day-to-day brightness variations.

## QUANTIFICATION AND STATISTICAL ANALYSIS

This work uses terminology from fluorescence microscopy. Here, fluorescence intensity refers to photon counts, which can be background-corrected (*I*) or -uncorrected (*S*),. Count rate (*CR*) denotes the frequency of photon counts and is generally calculated as 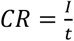 Each analysis might include different data points (e.g., some parameters are calculated from total photon emission, while others are calculated only from donor photon emission). These specifications are mentioned in each case.

As this is interdisciplinary work, the same parameters may be represented differently across analyses. Our intention is for these variables to be independently understood within each context.

These particularities are mentioned accordingly, always referring to the definitions presented in the above paragraph.

### Dimerization and DNA binding assays

Steady-state fluorescence anisotropy (*r*_*ss*_) was calculated from polarized fluorescence measurements according to **Equation 1**. Here, *F*_*p*_ and *F*_*s*_ represent the background-corrected fluorescence count rate, detected parallel or perpendicular to the excitation polarization, respectively, and *G* is the instrumental correction factor determined from standard measurements.

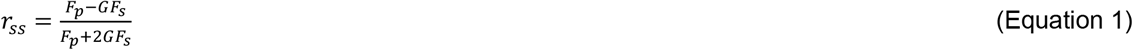

Anisotropy curves were fitted in OriginPro 2025b (OriginLab Corporation, Northampton, MA, United States) using the Hill equation, with the Hill coefficient fixed to n=1 for the dimerization curve and free for the DNA-binding analysis.

### Confocal Microscopy: Localization assay

Confocal image stacks were deconvolved using the Deconvolution Wizard in Huygens Professional 24.10 (Scientific Volume Imaging, Hilversum, Netherlands). Background levels were adjusted individually for each sample, while the acuity parameter and other algorithm settings were kept at default values. An axial chromatic offset of the Hoechst channel and the eGFP/mCherry channels was detected and corrected using the Chromatic Aberration Correction Wizard in Huygens. For FoxP1 constructs lacking nuclear localization, this correction step was omitted. Two-dimensional images of selected optical slices were visualized using Fiji^69^.

### Pulsed Interleaved Excitation Fluorescence Lifetime Imaging Microscopy

PIE-FLIM analysis consisted of four main steps: (1) segmentation of cellular nuclei, (2) extraction of time-resolved fluorescence data, (3) calculation of pixel-based mean photon arrival times and total fluorescence intensities and (4) determination of fluorescence lifetimes and modeling of D-A distance distributions from photon arrival histograms integrated over all pixels within a ROI to determine the fraction of FRET-active molecules. These steps are detailed below.

In polarization-resolved PIE-FLIM experiments, six fluorescence detection channels were recorded, differing in polarization (either parallel, p, or perpendicular, s, to the excitation), spectral detection window (green or red, corresponding to donor or acceptor emission, respectively), and excitation wavelength (green (D) or red (A), also referred to as prompt or delay time windows). These channels were grouped into three detection classes: (i) directly excited donor emission, (ii) FRET-sensitized acceptor emission, and (iii) directly excited acceptor emission. The first detection class corresponds to donor fluorescence following green excitation. Accordingly, FRET-sensitized acceptor emission follows excitation with the green light, and directly excited acceptor emission is detected in the delay time window following red excitation. Here, background-uncorrected fluorescence intensities are represented by the variables 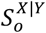, where the subscripts determine the detection orientation, the first superscript denotes the spectral detection window, and the second denotes the excitation wavelength.

#### Exporting intensity images

Intensity images for each group were generated by summing photons across frames using custom scripts in tttrlib^70^ and scikit-image^71^ and saved as 16-bit TIFF files.

#### Image segmentation

Nuclear regions of interest (ROIs) were defined based on total intensity across all channels. Non-nuclear FoxP1 variants were manually segmented in Fiji, producing binary masks (0 = background, 1 = ROI). Nuclear variants were segmented using a CellPose model^72,73^ trained on manually annotated masks. The CellPose-generated masks were verified for accuracy in Fiji.

#### Pixel-based mean photon arrival times

Mean photon arrival times for donor emission were computed per pixel using tttrlib and scikit-image, excluding pixels with < 20 photons, and exported as 32-bit floating-point TIFF files.

#### Subnuclear segmentation

Nuclei were subdivided into five sub-ROIs to determine local dimerization using two approaches: intensity-based and number-and-brightness (N&B). For the intensity-based segmentation, total background-uncorrected fluorescence intensity was calculated by summing the photon counts collected across all three detection classes described above 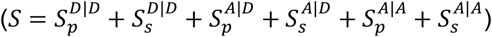. Subsequently, the Multi Otsu thresholding algorithm ^60^ implemented in scikit-image was used to partition each nucleus into “high”, “mid”, and “low” intensity regions. For the N&B-based segmentation, background-uncorrected fluorescence intensity fluctuations across the imaged frames were used to estimate the apparent number of molecules (*N*) and apparent brightness (*B*) per pixel^61,74^, as shown in **Equations 2** and **3**.

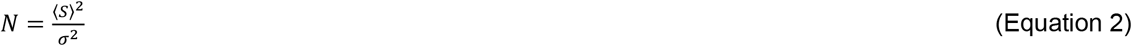

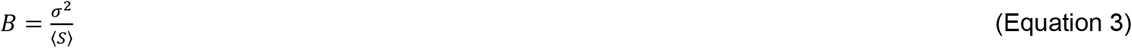

where *σ*^2^ represents the variance of the intensity distribution. Pixels were classified into “low-B” and “high-B” regions using an apparent brightness threshold of 1.5. All five regions were saved as 8-bit binary TIFF files for fluorescence decay export.

#### Fluorescence intensity decay and lifetime analysis

Photon arrival time histograms were constructed from the directly excited donor emission (bin size 20 ps) using tttrlib and scikit-image. Polarization artifacts introduced by high-NA objectives ^67,68^ and detection-sensitivity artifacts were corrected using correction factors and G-factors derived from fast- and slow-rotating fluorophores using ChiSurf ^75^. The average green correction factors obtained were *G* = 0.964, *l*_*p*_ = 0.059, and *l*_*s*_ = 0.305. Average donor lifetimes ⟨*τ*⟩_*x*_ were determined by iterative reconvolution of multi-exponential models (**Equation 4**) in *ChiSurf*.

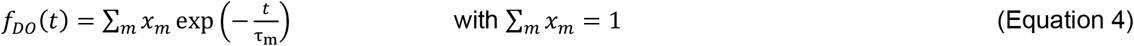

Briefly, experimental nuisances were introduced into the model by convolving with the IRF, computing partial polarization mixing, and scaling the models by the detectors’ detection sensitivities. The average fluorescence lifetime ⟨*τ*⟩_*x*_ (**Equation 5**) was calculated as species-weighted average.

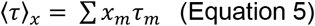

D-A distance distributions and the fractions of FRET-active molecules (*x*_*FRET*_) were determined by fitting a mixture model *f*(*t*) (**Equation 6**) to the data. This model includes the fluorescence decay of non-FRET molecules (*f*_*DO*_ (*t*)), and the fluorescence decay of FRET species,

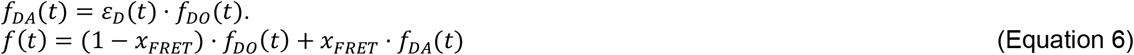

Here, *ε*_*D*_(*t*) is the FRET-induced donor decay that can be directly related to FRET rate constants and distances ^59^. Due to the flexibility of the C-terminal fluorescent protein tag, we related the FRET-induced donor decay to a Gaussian distance distribution (**Equation 7**), of half-width *σ*_*app*_ centered around 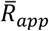. We computed the FRET-rate constants for distances for a single averaged orientation factor (*κ*^2^) of 2/3. Hence, recovered distances are apparent distances, *R*_*app*_.

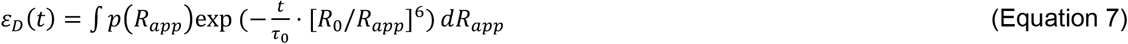

where the D-A distance distribution *p*(*R*_*app*_) is approximated by a normal distribution (**Equation 8**). A Förster radius *R*_0_ of 52 Å was used for the eGFP–mCherry FRET pair, consistent with published spectral overlap calculations for these fluorescent proteins ^76^.

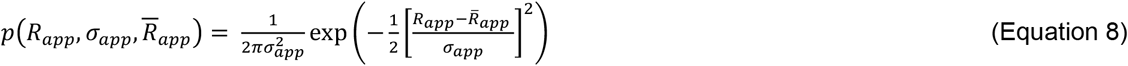

Analysis was performed in two steps across all nuclei. First, we fit the fluorescence decays of each individual cell, fixing *σ*_*app*_ to 6 Å, to find the central distance of the distribution. Second, fluorescence decays were fit again using the average values for 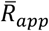 for each construct, with *σ*_*app*_ ranging from 5 to 20 Å, to probe *x*_*FRET*_. For subnuclear segmentation analysis, the five different sub-ROIs were fit jointly using the respective average 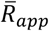 and a shared *σ*_*app*_. For FL R514H, two Gaussian populations were fit to account for multi-acceptor FRET. To avoid overparameterization and mixing of distance distributions, *σ*_*app*_ of the fast FRET-rate/short distance was fixed to 1 Å in both fitting rounds (with free and fixed 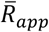, respectively). *Photon count rates and apparent stoichiometry determination*. Total photon count rates were calculated from all detection classes to estimate FoxP1 concentration, and were compensated for day-to-day variations in molecular brightness using calibration measurements. The ratio of donor to total protein concentration, known as apparent stoichiometry (*S*_*PIE,app*_) was calculated from these count rates as shown in **Equation 9**, following the superscript nomenclature described above.

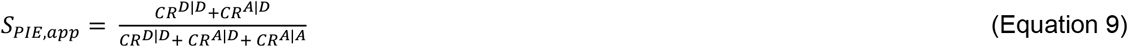

#### Correcting for local and relative FoxP1 concentration effects

To evaluate the dependence of *x*_*FRET*_ on FoxP1 protein concentration and on relative donor-to-acceptor ratio, we performed multiple linear regression analyses in OriginPro 2025b, using count rate and *S*_*PIE,app*_ as independent variables. To account for the identified dependencies, we constructed independent stoichiometry distributions for each D:A ratio by pooling data across all the studied FoxP1 constructs, and additionally generated a single *CR* distribution comprising all data points. Each stoichiometry distribution was modeled using a Gaussian normal function. To filter the data while ensuring at least 10 data points per experimental group, we retained nuclei with stoichiometry values within 1.5σ of the mean for subsequent analyses. This corresponded to ranges 0.435-0.945 for 1:1 D:A condition and 0.191-0.797 for the 1:5 condition. In contrast, the *CR* distribution followed a log-normal distribution; therefore, *CR* values were logarithmically transformed and fit to a Gaussian distribution. Nuclei with a *CR* value within 1σ of the mean were retained for further analyses, corresponding to a *CR* range of ∼240-1300 kHz. Box plots of *x*_*FRET*_ and ⟨*τ*⟩_*x*_ were generated with the filtered dataset. For visualization, the stoichiometries and *CR* from all ROIs were plotted as two-dimensional histograms both before and after application of the filtering criteria.

#### Nuclear distribution of FoxP1 homotypic interactions

To determine the nuclear distribution of FoxP1 homotypic interactions, we quantified the distribution of high-B pixels, which contain a higher proportion of oligomers and other higher-order configurations. First, we defined the nuclear periphery as a 0.5 µm rim based on histograms of the normalized radial distribution of high-B pixels for each construct and D:A condition. These histograms were generated by measuring the distance from each high-B pixel to the nearest point on the nuclear perimeter. From these radial distributions, we calculated the number of high-B in the rim and in the nuclear center and defined the high-B pixel (HB) enrichment score as shown in

**Equation 10**. Here, 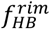 and 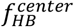represent the fraction of high-B pixels in the rim and in the center of the nucleus, respectively, which are calculated from the corresponding number of high-B pixels (*N*_*HB*_) relative to the total number of pixels (*N*) in each one of these regions. log_2_ calculation was taken to achieve a symmetric distribution around 0 for a uniform nuclear high-B pixel distribution.

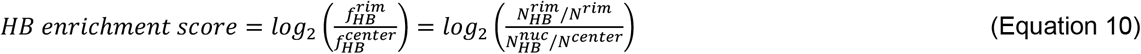

### Statistical analysis

All statistical analyses were performed in OriginPro 2025b. All distributional comparisons across groups are presented as box plots with box limits at the 25th and 75th percentiles, and whisker lengths determined by the outer data point that falls within 1.5 times the interquartile range above or below the 75th or 25^th^ percentile, respectively. The line in the box indicates the median, and the center of the box is the mean of the data distribution. Outliers were identified and removed independently from each of the groups in these distributional comparisons by calculating the maximum deviation from the mean using Grubb’s test at the 0.05 significance level. The distributional and statistical parameters shown in each box chart were calculated after outlier removal. All statistical differences were calculated by ANOVA. The number of symbols shown in all the plots denotes the level of statistical significance: one symbol indicates *p* < 0.05, two indicate *p* <0.01, and three indicate *p* < 0.001.

