## Supplementary material for "Multidomain Coupling Governs FoxP1 Assembly and Nuclear Compartmentalization": Document S1

[4] Current address: Biophysical Chemistry, Department of Chemistry and Chemical Biology, Technische Universität Dortmund, Dortmund, Germany

[6] Lead contact

### Supplementary Figures

#### Supplementary Figure 1

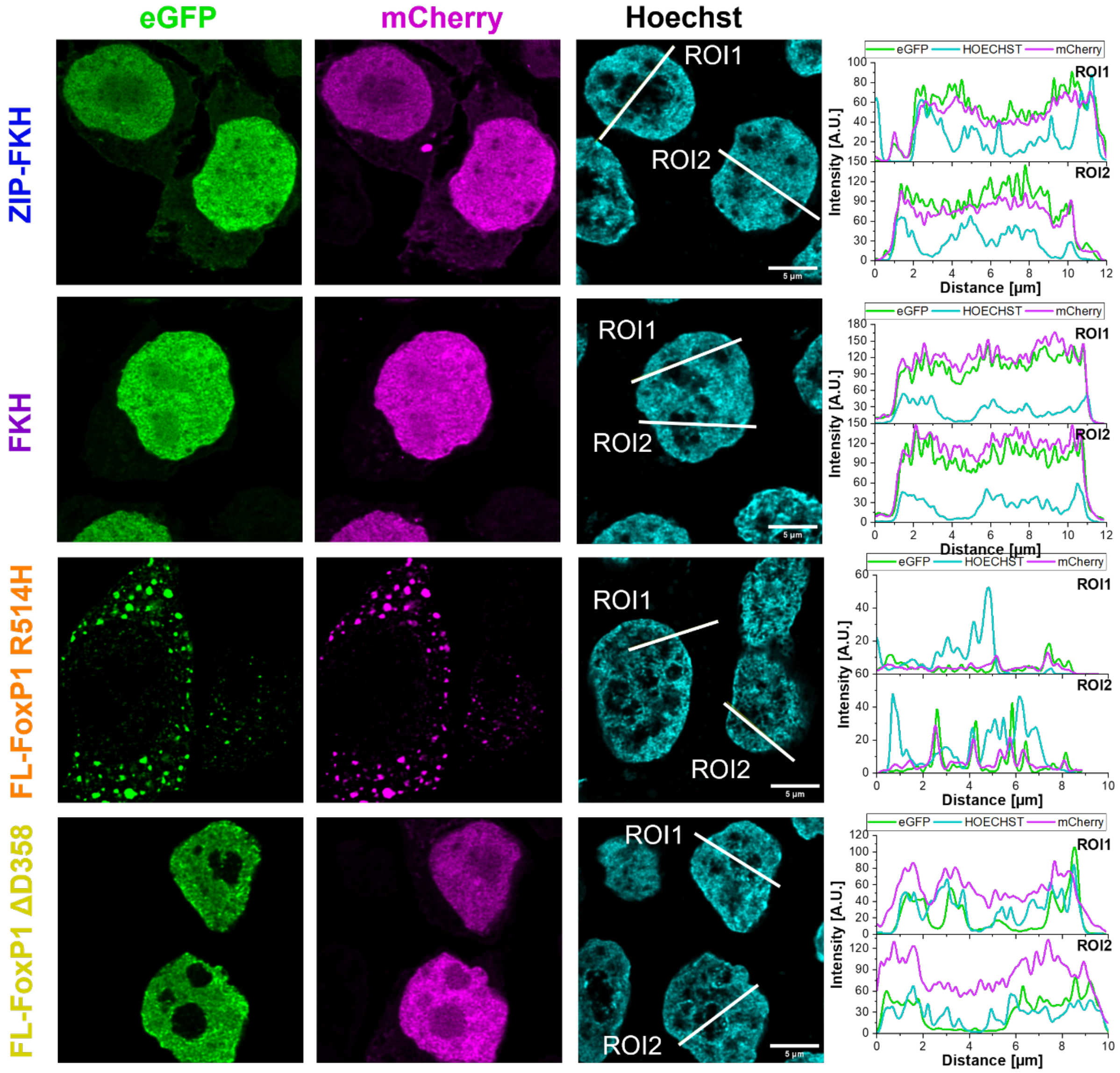

**Supplementary Figure 1. Representative confocal images of full-length and truncated FoxP1 variants.** Confocal image of a Hoechst-stained HEK293T cell nucleus co-transfected with eGFP- and mCherry-tagged FoxP1 constructs showing the expression of both constructs. Line profiles of the two regions of interest (ROIs) show the signal intensity in the three fluorescent channels. Scale bar: 5 µm.

#### Supplementary Figure 2

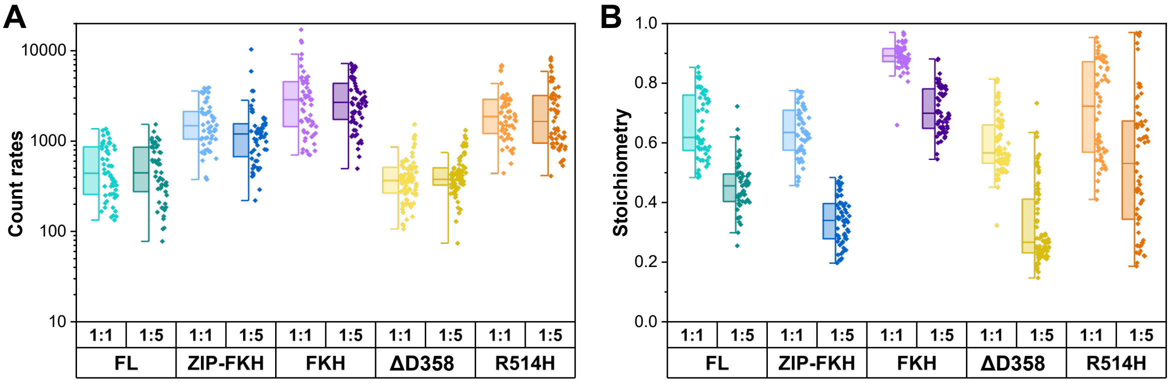

**Supplementary Figure 2. Total count rates and apparent Stoichiometries for all full nucleus ROIs. (A)** Total photon count rate computed as the sum of all count rates in the donor and acceptor channels after both donor and acceptor excitation. **(B)** The apparent Stoichiometry, $S_{PIE,app}$, was calculated as the ratio of the green (donor) and red (acceptor) signals in the prompt time window (i.e., donor excitation) to the total fluorescence intensity after donor and acceptor excitation. For details see Methods.

### Supplementary Figure 3

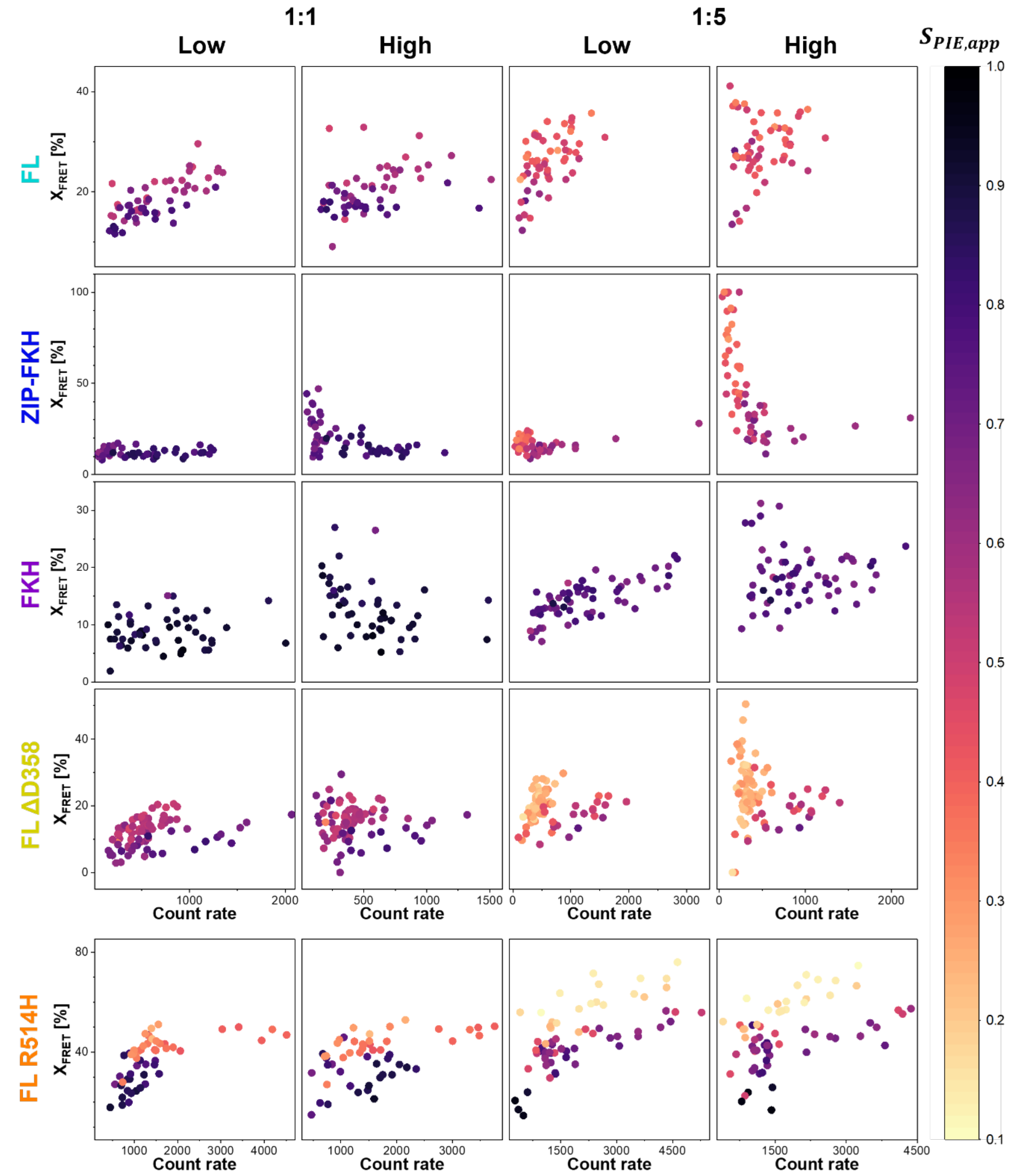

**Supplementary Figure 3. FRET characteristics of FoxP1 full-length and truncation variants depend on expression level and expression level ratio.** The fraction of molecules undergoing FRET ($x_{FRET}$) depends on stoichiometry ($S_{PIE,app}$), the expression level ratio of the donor relative to the total protein concentration, and the total photon count rate (CR), the protein expression level. $S_{PIE,app}$ is color-coded from yellow to dark violet in the range from 0.1 to 1.

#### Supplementary Figure 4

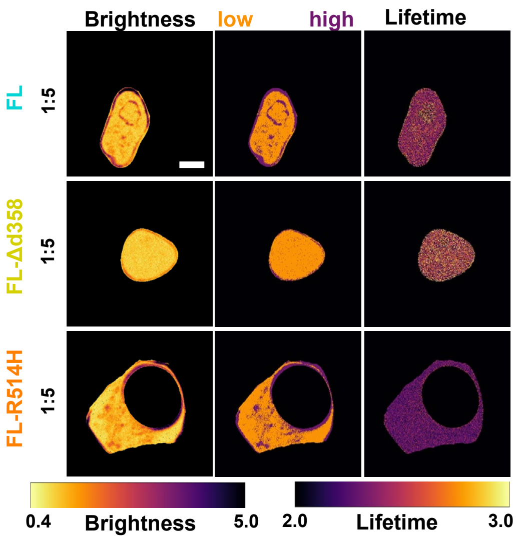

**Supplementary Figure 4. FL-FoxP1 homo-interaction of FL-FoxP1 variants.** Representative FL, FL ΔD358 and FL R514H nuclei (1:5), showing apparent brightness (left), nuclear subregions defined by N&B-based ROI segmentation (middle), and donor fluorescence lifetime per pixel (right). Scale bar: 5 µm.

#### Supplementary Figure 5

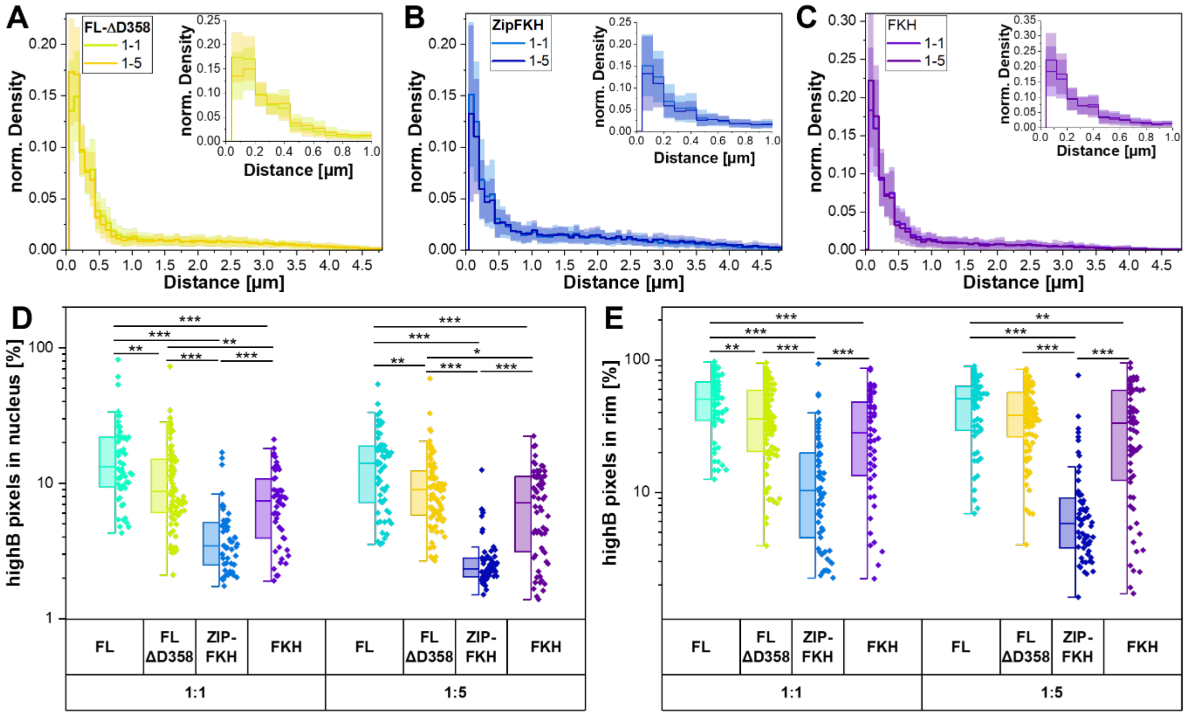

**Supplementary Figure 5. Distribution of high brightness pixels in the FoxP1-transfected nuclei. (A-C)** Normalized radial density of high-brightness pixels, where 0 µm denotes the nucleus border for **(A)** FoxP1 FL ΔD358, **(B)** ZIP-FKH and **(C)** FKH. **(D)** Fraction of high brightness pixels in the whole nucleus. **(E)** Fraction of high brightness pixels in the 0.5 µm rim.

#### Supplementary Figure 6

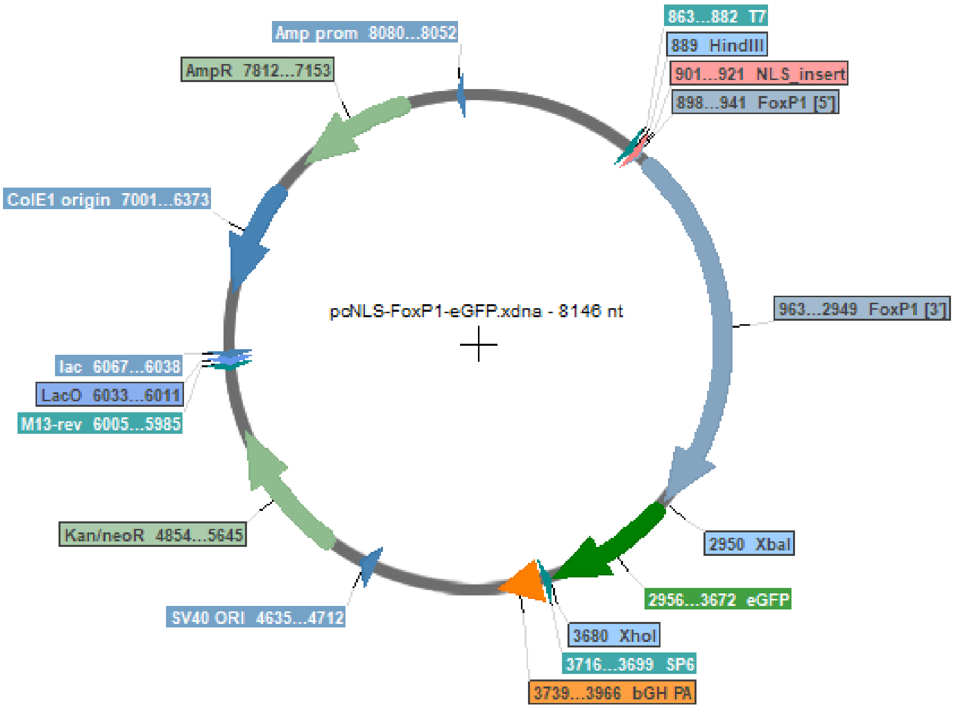

**Supplementary Figure 6. Plasmid map of full-length FoxP1-eGFP in pcDNA3.** Human full-length FoxP1 was synthesized and cloned into pcDNA between the HindIII and XbaI cleavage sites. N-terminally, the FoxP1-derived nuclear localization sequence RRRYSD was inserted. eGFP or mCherry was cloned C-terminally directly behind FoxP1 using the XbaI and XhoI cleavage sites without any additional linker sequences between the fluorescent protein and FoxP1.

### Supplementary Tables

#### Supplementary Table 1

**Supplementary Table 1. List of FoxP1 constructs generated.** The fluorescent proteins were added N- or C-terminally to FoxP1; in addition, a nuclear localization sequence (NLS) was tested to improve nuclear localization of the short fragment. RRRYSD is the NLS derived from FoxP1 (“FoxP1NLS”), while PKKRKV is derived from SV40 (“SV40NLS”). Constructs marked with * were used in this study.

| Construct name based on sequence | Functional modification to FoxP1 sequence | Localization |
|---|---|---|
| eGFP-FL FoxP1 | None | Whole cell |
| FL FoxP1-eGFP | None | Nucleus |
| SV40NLS-FL FoxP1-eGFP | None | Nucleus |
| FoxP1NLS-eGFP-FL FoxP1 | None | Cytosol |
| FoxP1NLS-FL FoxP1-eGFP* | None | Nucleus |
| SV40NLS-ZIP-FKH-eGFP | ZIP-FKH region (aa 345-677) only | Nucleus |
| SV40NLS-eGFP-ZIP-FKH | ZIP-FKH region (aa 345-677) only | Whole cell |
| FoxP1NLS-ZIP-FKH-eGFP* | ZIP-FKH region (aa 345-677) only | Whole cell |
| FoxP1NLS-FKH-eGFP* | FKH domain (aa 462-677) only | Nucleus |
| FL FoxP1-mCherry | None | Whole cell (weak expression) |
| mCherry-FL FoxP1 | None | Nucleus |
| SV40NLS-FL FoxP1-mCherry | None | Nucleus |
| FoxP1NLS-FL FoxP1-mCherry* | None | Nucleus |
| SV40NLS-ZIP-FKH-mCherry* | ZIP-FKH region (aa 345-677) only | Nucleus |
| FoxP1NLS-ZIP-FKH-mCherry | ZIP-FKH region (aa 345-677) only | Whole cell |
| SV40NLS-mCherry-ZIP-FKH* | ZIP-FKH region (aa 345-677) only | Whole cell |
| FoxP1NLS-FKH-mCherry* | FKH domain (aa 462-677) only | Nucleus |
| FoxP1NLS-FL FoxP1 R514H-eGFP* | R514H mutation | Partially nucleus/partially whole cell (puncta) |
| FoxP1NLS-FL FoxP1 ΔD358-eGFP* | ΔD358 deletion | Nucleus |
| FoxP1NLS-FL FoxP1 R514H-mCherry* | R514H mutation | Partially nucleus/partially whole cell (puncta) |
| FoxP1NLS-FL FoxP1 ΔD358-mCherry* | ΔD358 deletion | Nucleus |

#### Supplementary Table 2

**Supplementary Table 2. Correction factors for data filtering and comparability.** FRET characteristics of FoxP1 full-length and truncation variants depend on expression level (Count rate) and expression level ratio (Stoichiometry). The correlations shown in Supplementary Figure 3 were fitted with linear equations to obtain their slope (“Coefficient”). The p-value indicates the likelihood of deviations from a straight line with slope = 0. $R^2$ denotes the goodness of the fit.

| ***D:A ratio*** | ***Construct*** | ***Bright-ness*** | ***Z- intercept*** | | ***Stoichiometry dependence*** | | ***Count rate dependence*** | | $R^2$ |
|---|---|---|---|---|---|---|---|---|---|
| | | | Coefficient | P-value | Coefficient | P-value | Coefficient | P-value | |
| 1:1 | FL | Low | 0.24±0.02 | 1.5E-16 | -0.16±0.03 | 2.9E-7 | 6.9E-5±8.7E-6 | 1.1E-10 | 0.71 |
| | | High | 0.33±0.03 | 3.8E-13 | -0.22±0.05 | 1.5E-5 | 4.0E-5±1.6E-5 | 0.017 | 0.38 |
| | ZIP-FKH | Low | 0.22±0.03 | 4.7E-8 | -0.13±0.05 | 7.8E-3 | 1.2E-5±9.3E-6 | 0.19 | 0.092 |
| | | High | 0.47±0.14 | 10.0E-4 | -0.29±0.19 | 0.13 | -1.4E-4±4.6E-5 | 4.5E-3 | 0.32 |
| | FKH | low | 0.37±0.07 | 4.0E-6 | -0.32±0.08 | 1.9E-4 | 7.1E-6±8.5E-6 | .41 | 0.23 |
| | | High | 0.72±0.11 | 2.1E-8 | -0.62±0.12 | 3.5E-6 | -5.6E-5±1.8E-5 | 3.0E-3 | 0.44 |

| | | | | | | | | | |
|---|---|---|---|---|---|---|---|---|---|
| | FL ΔD358 | Low | 0.27±0.02 | 1.2E-20 | -0.30±0.04 | 2.2E-12 | 6.7E-5± 9.9E-6 | 2.1E-9 | 0.49 |
| | | High | 0.27±0.03 | 1.5E-11 | -0.21±0.06 | 5.0E-4 | 2.2E-5± 2.4E-5 | 0.37 | 0.12 |
| | FL R514H | Low | 0.46±0.03 | 2.0E-22 | -0.24±0.03 | 4.9E-10 | 3.4E-5± 8.7E-6 | 3.0E-4 | 0.70 |
| | | High | 0.45±0.03 | 4.2E-22 | -0.24±0.03 | 4.9E-10 | 4.3E-5± 9.7E-6 | 6.0E-5 | 0.64 |
| 1:5 | FL | Low | 0.34±0.03 | 5.6E-15 | -0.28±0.06 | 3.6E-5 | 7.2E-5± 1.5E-5 | 1.1E-5 | 0.49 |
| | | High | 0.46±0.06 | 5.8E-11 | -0.34±0.10 | 1.5E-3 | -6.4E-6± 3.2E-5 | 0.84 | 0.15 |
| | ZIP-FKH | Low | 0.27±0.02 | 6.6E-19 | -0.25±0.04 | 3.1E-7 | 4.4E-5± 9.5E-6 | 1.8E-5 | 0.39 |
| | | High | 1.23±0.13 | 1.2E-13 | -1.4±0.28 | 5.7E-6 | -1.6E-4± 8.4E-5 | 6.9E-2 | 0.47 |
| | FKH | low | 6.6E-2± 3.3E-2 | 4.8E-2 | 5.2E-2± 4.4E-2 | 0.24 | 3.3E-5± 4.1E-6 | 3.3E-11 | 0.51 |
| | | high | 0.25±0.07 | 9.9E-4 | -9.2E-2± 9.5E-2 | 0.34 | 1.4E-6± 1.3E-5 | 0.91 | -1.7 E-2 |
| | FL ΔD358 | Low | 0.26±0.01 | 9.7E-41 | -0.33±0.04 | 4.0E-14 | 8.1E-5± 1.2E-5 | 2.5E-9 | 0.52 |
| | | High | 0.31±0.03 | 6.1E-20 | -0.30±0.10 | 4.7E-3 | 4.8E-5± 4.7E-5 | 0.31 | 0.10 |
| | FL R514H | Low | 0.53±0.02 | 5.2E-37 | -0.33±0.02 | 1.4E-20 | 4.7E-5± 4.9E-6 | 9.5E-14 | 0.86 |
| | | High | 0.55±0.02 | 3.7E-35 | -0.36±0.03 | 1.5E-19 | 5.2E-5± 7.5E-6 | 2.5E-9 | 0.80 |

### Supplementary Table 3

**Supplementary Table 3. Full-length FoxP1 amino acid sequence**. Dark blue RRRYSD is the nuclear-localization sequence derived from FoxP1 and inserted in front of all constructs. Dark grey indicates the Q-rich region, salmon the Zinc-finger domain, light blue the leucine zipper domain, and purple the Forkhead DNA-binding domain. The aspartic acid highlighted in yellow marks the deleted ΔD358, and the arginine highlighted in red marks the mutation R514H.

```
RRRYSD
        10         20         30         40         50
MMQESGTETK SNGSAIQNGS GGSNHLLECG GLREGRSNGE TPAVDIGAAD
        60         70         80         90        100
LAHAQQQQQQ ALQVARQLLL QQQQQQQVSG LKSPKRNDKQ PALQVPVSVA
       110        120        130        140        150
MMTPQVITPQ QMQQILQQQV LSPQQLQVLL QQQQALMLQQ QQLQEFYKKQ
       160        170        180        190        200
QEQLQLQLLQ QQHAGKQPKE QQQVATQQLA FQQQLLQMQQ LQQQHLLSLQ
       210        220        230        240        250
RQGLLTIQPG QPALPLQPLA QGMIPTELQQ LWKEVTSAHT AEETTGNNHS
       260        270        280        290        300
SLDLTTTCVS SSAPSKTSLI MNPHASTNGQ LSVHTPKRES LSHEEHPHSH
       310        320        330        340        350
PLYGHGVCKW PGCEAVCEDF QSFLKHLNSE HALDDRSTAQ CRVQMQVVQQ
       360        370        380        390        400
LELQLAKDKE RLQAMMTHLH VKSTEPKAAP QPLNLVSSVT LSKSASEASP
       410        420        430        440        450
QSLPHTPTTP TAPLTPVTQG PSVITTTSMH TVGPIRRRYS DKYNVPISSA
       460        470        480        490        500
DIAQNQEFYK NAEVRPPFTY ASLIRQAILE SPEKQLTLNE IYNWFTRMFA
       510        520        530        540        550
YFRRNAATWK NAVRHNLSLH KCFVRVENVK GAVWTVDEVE FQKRRPQKIS
       560        570        580        590        600
GNPSLIKNMQ SSHAYCTPLN AALQASMAEN SIPLYTTASM GNPTLGNLAS
```

```
       610        620        630        640        650
AIREELNGAM EHTNSNESDS SPGRSPMQAV HPVHVKEEPL DPEEAEGPLS
       660        670
LVTTANHSPD FDHDRDYEDE PVNEDME
```

#### Supplementary Table 4

**Supplementary Table 4. Full-length *FoxP1* gene sequence**. Italic bold bases indicate HindIII and XbaI cloning sites. Color code is identical to the protein sequence shown in Supplementary Table 3.

| | |
|---|---|
| 1 | ***AAG CTT*** AAC ATG **CGC CGC CGC TAT AGC GAT** ATG CAG GAA AGC GGC ACC GAA |
| 55 | ACC AAA AGC AAC GGC AGC GCG ATT CAG AAC GGC AGC GGC GGC AGC AAC CAT |
| 109 | CTG CTG GAA TGC GGC GGC CTG CGC GAA GGC CGC AGC AAC GGC GAA ACC CCG |
| 163 | GCG GTG GAT ATT GGC GCG GCG GAT CTG GCG CAT GCG **CAG CAG CAG CAG CAG** |
| 217 | **CAG GCG CTG CAG GTG GCG CGC CAG CTG CTG CTG CAG CAG CAG CAG CAG CAG** |
| 271 | **CAG GTG AGC GGC CTG AAA AGC CCG AAA CGC AAC GAT AAA CAG CCG GCG CTG** |
| 325 | **CAG GTG CCG GTG AGC GTG GCG ATG ATG ACC CCG CAG GTG ATT ACC CCG CAG** |
| 379 | **CAG ATG CAG CAG ATT CTG CAG CAG CAG GTG CTG AGC CCG CAG CAG CTG CAG** |
| 433 | **GTG CTG CTG CAG CAG CAG CAG GCG CTG ATG CTG CAG CAG CAG CAG CTG CAG** |
| 587 | **GAA TTT TAT AAA AAA CAG CAG GAA CAG CTG CAG CTG CAG CTG CTG CAG CAG** |
| 541 | **CAG CAT GCG GGC AAA CAG CCG AAA GAA CAG CAG CAG GTG GCG ACC CAG CAG** |
| 595 | **CTG GCG TTT CAG CAG CAG CTG CTG CAG ATG CAG CAG CTG CAG CAG CAG CAT** |
| 649 | **CTG CTG AGC CTG CAG CGC CAG GGC CTG CTG ACC ATT CAG CCG GGC CAG CCG** |
| 703 | **GCG CTG CCG CTG CAG CCG CTG GCG CAG GGC ATG ATT CCG ACC GAA CTG CAG** |
| 757 | **CAG** CTG TGG AAA GAA GTG ACC AGC GCG CAT ACC GCG GAA GAA ACC ACC GGC |
| 811 | AAC AAC CAT AGC AGC CTG GAT CTG ACC ACC ACC TGC GTG AGC AGC AGC GCG |
| 865 | CCG AGC AAA ACC AGC CTG ATT ATG AAC CCG CAT GCG AGC ACC AAC GGC CAG |
| 919 | CTG AGC GTG CAT ACC CCG AAA CGC GAA AGC CTG AGC CAT GAA GAA CAT CCG |
| 973 | CAT AGC CAT CCG CTG TAT GGC CAT **GGC GTG TGC AAA TGG CCG GGC TGC GAA** |
| 1027 | **GCG GTG TGC GAA GAT TTT CAG AGC TTT CTG AAA CAT CTG AAC AGC GAA CAT** |
| 1081 | GCG CTG GAT GAT CGC AGC ACC GCG CAG TGC CGC GTG CAG **ATG CAG GTG GTG** |
| 1135 | **CAG CAG CTG GAA CTG CAG CTG GCG AAA GAT AAA GAA CGC CTG CAG GCG ATG** |
| 1189 | **ATG ACC CAT** CTG CAT GTG AAA AGC ACC GAA CCG AAA GCG GCG CCG CAG CCG |
| 1243 | CTG AAC CTG GTG AGC AGC GTG ACC CTG AGC AAA AGC GCG AGC GAA GCG AGC |
| 1297 | CCG CAG AGC CTG CCG CAT ACC CCG ACC ACC CCG ACC GCG CCG CTG ACC CCG |
| 1351 | GTG ACC CAG GGC CCG AGC GTG ATT ACC ACC ACC AGC ATG CAT ACC GTG GGC |
| 1405 | CCG ATT **CGC CGC CGC TAT AGC GAT** AAA TAT AAC GTG CCG ATT AGC AGC GCG |
| 1459 | GAT ATT GCG CAG AAC CAG GAA TTT TAT AAA AAC **GCG GAA GTG CGC CCG CCG** |
| 1513 | **TTT ACC TAT GCG AGC CTG ATT CGC CAG GCG ATT CTG GAA AGC CCG GAA AAA** |
| 1567 | **CAG CTG ACC CTG AAC GAA ATT TAT AAC TGG TTT ACC CGC ATG TTT GCG TAT** |
| 1621 | **TTT CGC CGC AAC GCG GCG ACC TGG AAA AAC GCG GTG CGC CAT AAC CTG AGC** |
| 1675 | **CTG CAT AAA TGC TTT GTG CGC GTG GAA AAC GTG AAA GGC GCG GTG TGG ACC** |
| 1729 | **GTG GAT GAA GTG GAA TTT CAG AAA CGC CGC CCG CAG AAA** ATT AGC GGC AAC |
| 1783 | CCG AGC CTG ATT AAA AAC ATG CAG AGC AGC CAT GCG TAT TGC ACC CCG CTG |
| 1837 | AAC GCG GCG CTG CAG GCG AGC ATG GCG GAA AAC AGC ATT CCG CTG TAT ACC |
| 1891 | ACC GCG AGC ATG GGC AAC CCG ACC CTG GGC AAC CTG GCG AGC GCG ATT CGC |
| 1945 | GAA GAA CTG AAC GGC GCG ATG GAA CAT ACC AAC AGC AAC GAA AGC GAT AGC |
| 1999 | AGC CCG GGC CGC AGC CCG ATG CAG GCG GTG CAT CCG GTG CAT GTG AAA GAA |
| 2053 | GAA CCG CTG GAT CCG GAA GAA GCG GAA GGC CCG CTG AGC CTG GTG ACC ACC |
| 2107 | GCG AAC CAT AGC CCG GAT TTT GAT CAT GAT CGC GAT TAT GAA GAT GAA CCG |
| 2161 | GTG AAC GAA GAT ATG GAA ***TCT AGA*** |
